# RNA-binding proteins activate transcription through defined molecular grammars

**DOI:** 10.64898/2026.05.29.728737

**Authors:** Haoran Wang, Lydia Phillips, Dervla Moore-Frederick, Bailey K. Webster, Deyuan Xu, Sid Srivastava, Grant Mowry, Emily Wang, Arth Banka, Changfeng Deng, David M. Chenoweth, Jonathan E. Henninger

## Abstract

Transcription factors (TFs) regulate gene expression through interactions with DNA, RNA, and proteins. RNA-binding proteins (RBPs) also assemble near regulatory elements and mediate RNA processing, yet their perturbation causes transcriptional defects. Here, we find select RBPs activate transcription through latent activation domains akin to TFs. RBP activators regulate distinct genes and interact with transcriptional condensates. Their activation domains are enriched in aromatic and polar residues but depleted of basic residues – essential features that are conserved and partially mimic TF activation domains. We validated additional RBP activators across the human proteome based on this molecular grammar, including the C-terminal domain (CTD) of RPB1, the catalytic subunit of RNA polymerase II. RPB1-CTD activates transcription by recruiting coactivators, demonstrating a non-enzymatic function in transcriptional regulation. These findings position RBPs and RPB1 as transcriptional regulators, explain coupling between transcription and RNA processing, and reveal RNA-RBP regulatory networks that parallel DNA-TF networks.

## Introduction

Over 15% of human protein-coding genes encode proteins that bind to nucleic acid. These include transcription factors (TFs) that bind DNA molecules to regulate transcription initiation and elongation^1^, as well as RNA-binding proteins (RBPs) that bind nuclear and cytoplasmic RNA molecules to regulate post-transcriptional processes^2^. Although canonical models of gene expression typically sort DNA- and RNA-binding factors to distinct functional classes^3,4^, multiple lines of evidence point to a significant overlap in their binding preferences and function^5,6^. For instance, TFIIIA, the first TF discovered with C_2_H_2_ zinc-finger domains in *Xenopus laevis*, binds both DNA and RNA^7,8^. TFIIIA binds to DNA promoter elements in the 5S ribosomal RNA gene to regulate transcription, yet it also binds to newly synthesized 5S pre-ribosomal RNA to regulate 5S RNA transport^9^. More recent studies have revealed that a large fraction of TFs bind to RNA molecules^10^, and in some cases, regulate post-transcriptional processes^11–14^. In a similar fashion, some proteins with structured RNA-binding domains are capable of binding DNA^5,15–17^. Perturbation of a few RBPs, including those involved in RNA splicing, can also dysregulate RNA polymerase II (RNAPII) processivity and transcription^18–20^. Together, these studies inspire a more integrated model for the control of gene expression that incorporates interactions and feedback between TFs, RBPs, DNA, and RNA^5,21^.

TFs function by binding to cis-regulatory elements and recruiting transcription cofactors. Such binding events are separately encoded in specific binding domains, including DNA-binding domains that selectively target DNA motifs and effector domains that recruit co-activators or co-repressor proteins^1^. Co-activators and co-repressors alter DNA accessibility and the chemical properties of chromatin, and in some cases, directly recruit RNAPII or members of the pre-initiation complex^22,23^. Whereas the DNA-binding domains retain stereotypic structures, TF effector domains are more structurally diverse and often overlap intrinsically disordered regions (IDRs)^24^. These IDRs are thought to form multivalent, specific interactions with other transcriptional proteins, and in some cases, are sufficient to maintain proper TF genomic localization even in the absence of a DNA-binding domain^25^. Specific amino acid enrichments within TF IDRs, including leucine, aromatic and acidic amino acids, contribute to their ability to interact with coactivators and mediate transcriptional regulation^26–28^. A sufficiently high density of TFs, cofactors, and RNAPII can drive the formation of transcriptional condensates, which are dynamic network fluids, near genes^29–31^. The contribution of site-specific binding to nucleic acids and compatible molecular interactions between protein partners together control the activation of genes.

RBPs regulate gene expression at multiple levels, from co-transcriptional RNA processing to post-transcriptional RNA metabolism. In the nucleus, RBPs can bind to nascently transcribed RNAs to either directly process the RNA, as is the case for splicing factors, or to regulate such processes, as is the case for RBPs such as RNA-Binding Motif (RBM) family proteins, heterogeneous nuclear ribonucleoproteins (HNRNPs), and serine/arginine-rich (SR) proteins^32^. RBPs also bind to regulatory non-coding RNAs and shape local genome structure^33,34^. Similar to TFs, RBPs have distinct domains that recognize nucleic acid and large IDRs that enable multivalent protein-protein interactions. Disease-associated mutations in RBPs often alter RNA-binding specificity and impair splicing fidelity^35^, but emerging evidence indicates that these mutations can also perturb transcription^18–20^. Indeed, both transcription and co-transcriptional RNA processing are coupled processes in close proximity. For instance, during transcription, progressive phosphorylation of the RPB1-CTD, the catalytic subunit of RNAPII, helps recruit and assemble the spliceosome^36^. Conversely, RBPs can enhance RNAPII elongation rates, and the presence of introns increases nascent transcription. These observations raise an unresolved question: do RBPs influence transcription indirectly through RNA processing, or can they act directly as regulators of transcription itself?

Here, we provide evidence that select RBPs act as direct transcriptional activators beyond their role in co-transcriptional RNA processing. RBP activators have IDRs that function similar to TF activation domains. Acute RBP overexpression can regulate endogenous gene transcription of specific genes without impacting their splicing. RBP activators interact with RNAPII or transcription coactivators, like MED1, a subunit of the Mediator coactivator complex. Protein sequence analysis of RBP IDRs revealed an enriched molecular grammar that is necessary for transcription activation and predictive of proteome-wide RBP activators. Through molecular grammar enrichments, we find that the RPB1-CTD can recruit coactivators to activate gene transcription, suggesting a non-enzymatic function for RNAPII in transcription. Together, these findings reveal a non-canonical role for RBPs in transcription regulation and help explain the coupling between transcription and co-transcriptional RNA processing.

## Results

### Select chromatin-enriched RBPs stimulate transcription of a reporter gene

To enable gene-specific transcriptional regulation, TFs bind to chromatin at enhancers and promoters. Similar to TFs, select chromatin-enriched RBPs also assemble near gene regulatory elements^37^. For example, RBPs like FUS, TDP43, RBM proteins, and HNRNPs enrich at enhancers and promoters similar to the TF TAL1 near the *FOSL1* gene in human K562 leukemia cells (Figure 1A). This enrichment is observed across multiple enhancers, clusters of enhancers called super-enhancers, and promoters in these cells (Figure S1A). Although these factors have mainly been understood to regulate post-transcriptional RNA processing, perturbation of a few RBPs has been shown to disrupt transcriptional processes, suggestive of dual functions^18–20^.

**Figure 1.**
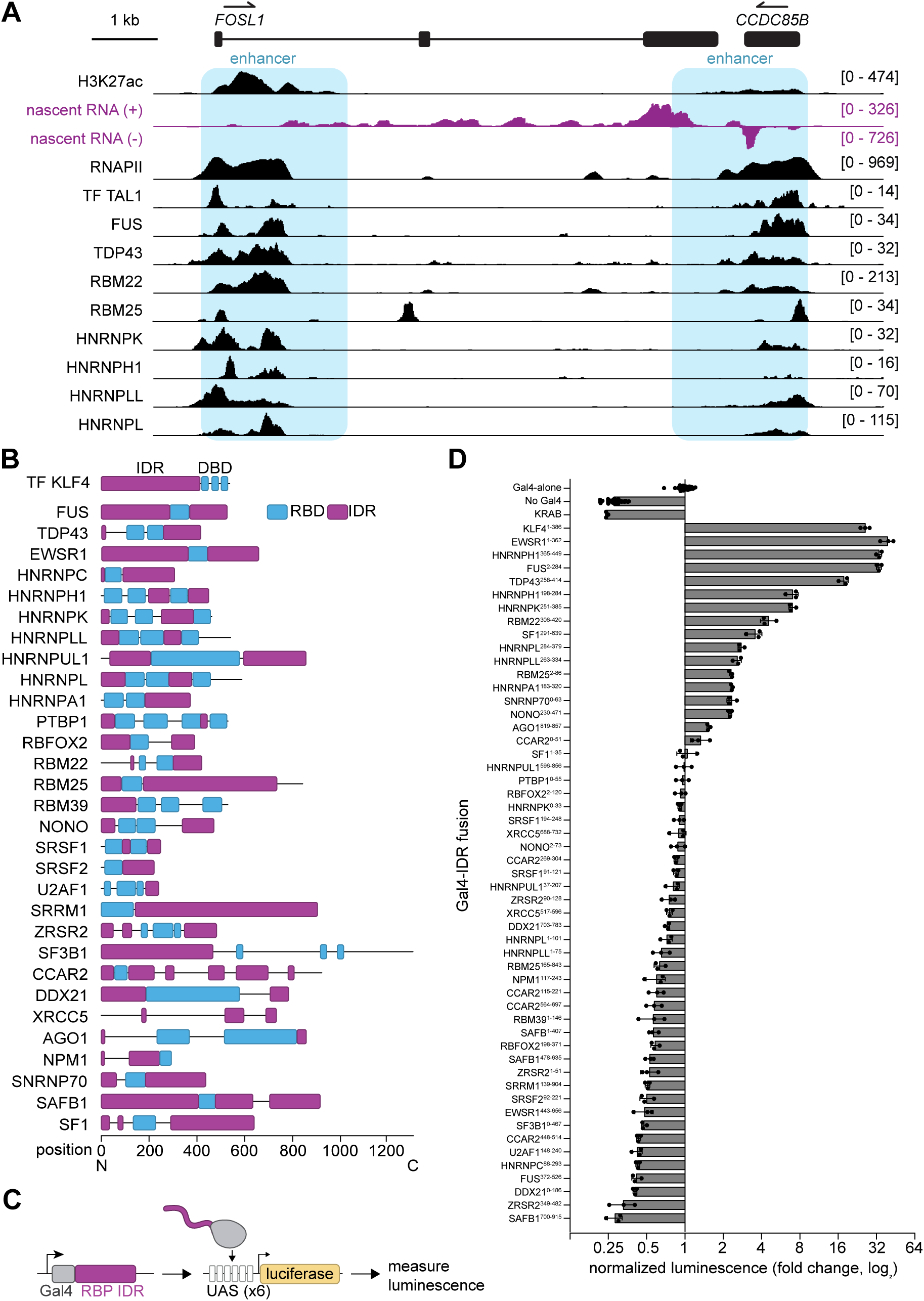
Select chromatin-enriched RBPs stimulate transcription of a reporter gene. A. TT-seq signal and ChIP-seq signal of H3K27ac, RNA Pol II, and select RBPs at the *FOSL1* locus in human K562 cells. B. Breakdown of RNA-binding domains and IDRs in select RBPs. C. Experimental scheme for Gal4-RBP IDR luciferase reporter assay. D. Bar plots showing the normalized luminescence of Gal4-RBP IDR luciferase assay. Values are normalized to the Gal4-alone condition and displayed on a log_2_ scale (statistics in Table S1).

We wondered if a subset of chromatin-enriched RBPs could function like TFs to directly regulate transcriptional output. TFs typically possess structured DNA-binding domains and intrinsically disordered effector domains that activate or repress transcription through cofactor recruitment^1,38^. Chromatin-enriched RBPs also have structured nucleic acid binding domains and IDRs^39^ (Figure 1B). We tested whether these regions had the potential to activate transcription using a classic transcriptional reporter assay^40^ that has been broadly successful in identifying TF activation domains^41^. We selected 30 chromatin-enriched RBPs, fused their IDRs (n = 52) to the heterologous DNA-binding domain (DBD) of Gal4 from yeast, and co-transfected expression plasmids with a luciferase reporter in human cells (Figure 1C). Several RBP IDRs were capable of activating reporter expression significantly above baseline levels within 4 hours of transfection (Figure 1D). These “RBP activators” exhibited a range of activation potentials, some of which surpassed the activation potential of the TF KLF4. Fusion Gal4-RBP-IDRs were expressed at similar levels in cells, suggesting that reporter expression reflected intrinsic activation potential (Figure S1B). RBP IDRs that exhibited activation potential similar to Gal4-DBD alone or to a repressive domain (Gal4-KRAB) were labeled as “non-activators” instead of repressors, which was confirmed by an alternative reporter assay using a stronger promoter (Figure S1C). The reporter is intronless, and RBP-mediated differences in luminescence were concordant with luciferase mRNA levels, suggesting that observed differences in luminescence were due to transcriptional rather than post-transcriptional effects (Figure S1D, S1E). RBP activators included several proteins that have been implicated in transcriptional control, such as EWSR1, TDP43, and FUS^16,42–45^. The assay also revealed many other RBP activators, including known RNA splicing regulator HNRNPs (e.g. HNRNPH1, HNRNPLL, HNRNPL, HNRNPA1), RBM proteins (RBM22 and RBM25), SF1, SNRNP70, and NONO. Together, these results show that select RBP IDRs are sufficient to activate transcription and suggest that these proteins could play functional roles beyond co-transcriptional RNA splicing.

### RBP activators regulate expression of endogenous genes

TFs regulate distinct sets of genes, including genes associated with cell identity. Because RBP IDRs have the potential to stimulate gene transcription (Figure 1), we next sought to determine if RBP activators regulate endogenous genes. We generated stable human K562 cell lines capable of inducible expression of Halo-tagged RBP activators SNRNP70, HNRNPA1, FUS, RBM22, and HNRNPH1, and acutely induced expression of these factors for 4 hours to potentially enrich for transcriptional, rather than post-transcriptional, effects on gene expression. Dozens of genes were differentially expressed when comparing RNA sequencing between RBP activators and a control cell line expressing Halo tag alone (Figure 2A, Figure S2A-E, Table S2). Interestingly, each RBP activator tended to stimulate distinct sets of genes (Figure 2A), a feature commonly observed for cell-specific TFs. Most differentially regulated genes were RBP-specific, but some factors, such as FUS and RBM22 or HNRNPA1 and SNRNP70, shared some differentially expressed genes in common (Figure 2B, Figure S2F). Few differentially regulated genes were co-regulated by multiple RBPs, suggesting that RBP-mediated activation or repression is more gene-specific than generic.

**Figure 2.**
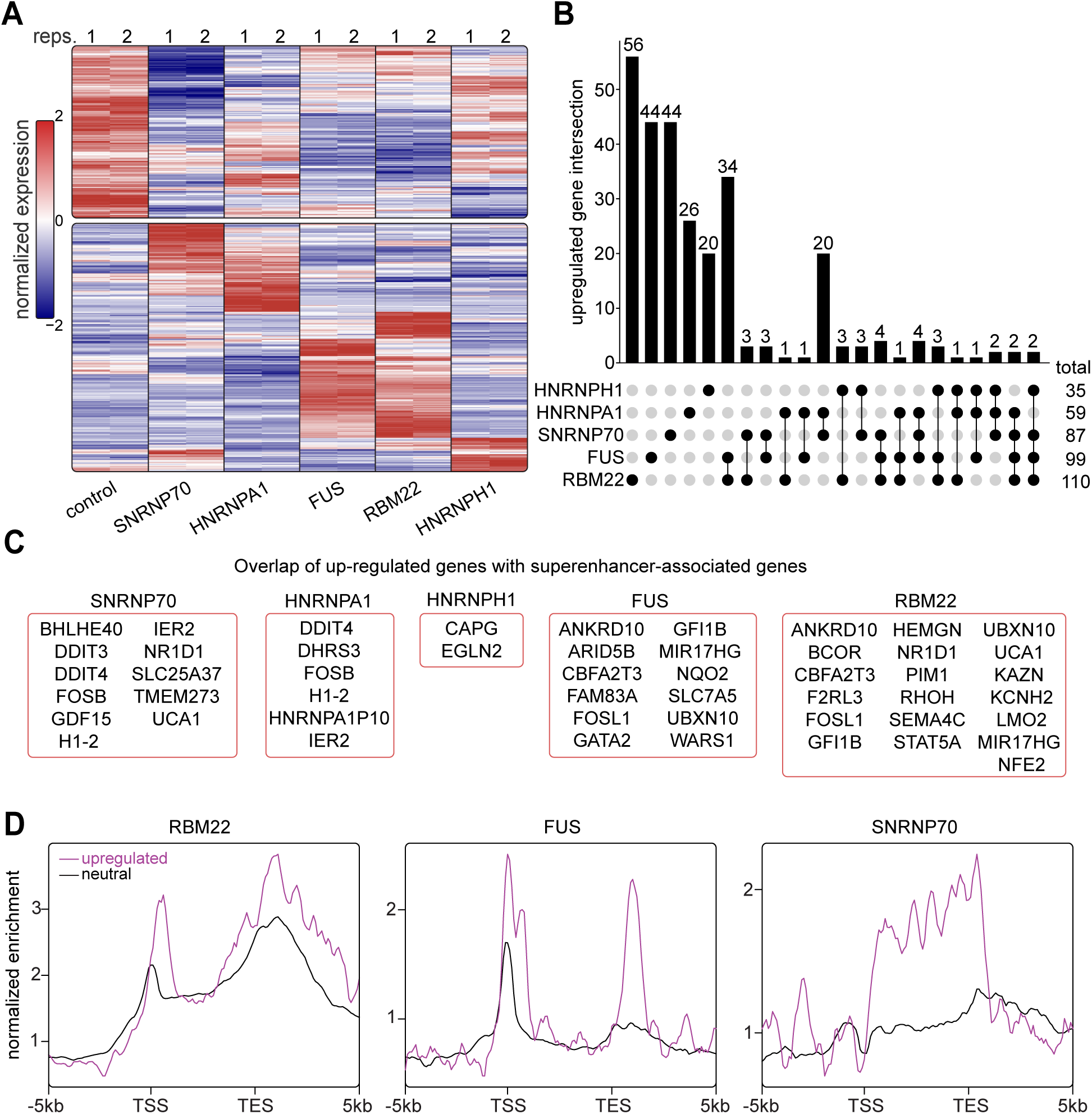
RBP activators regulate expression of endogenous genes. A. Heatmap of differentially expressed genes in doxycycline-inducible RBP overexpression K562 cell lines after 4 hours of treatment. Both the downregulated (top) and upregulated (bottom) genes compared to control are grouped together. B. Upset plot showing the upregulated gene intersection between 5 RBPs. C. Overlap of upregulated genes of specific RBPs with super-enhancer-associated genes. D. RBP ChIP-seq profile plots using upregulated and neutral super-enhancer-associated genes upon RBP overexpression.

Among the 1600 TFs^1^, many TFs are expressed in a cell-type specific fashion and regulate sets of genes associated with cell-identity. Such genes can be regulated by clusters of coordinating enhancers called super-enhancers, as these genes require selective but stable expression in cognate cell types^46,47^. Analysis of RNA-sequencing upon acute expression of RBP activators revealed that, similar to TFs, RBP activators regulate distinct super-enhancer associated genes (Figure 2C). Since RBP activators enrich at gene regulatory elements in proximity to super-enhancer associated genes, as is the case with FOSL1 (Figure 1A), we wondered if native chromatin occupancy was higher at activated compared to non-activated genes. Indeed, chromatin occupancy of endogenous RBPs at activated super-enhancer associated genes was higher compared to their non-activated counterparts, which would be expected if RBPs were regulating transcriptional processes (Figure 2D). Changes in RNA levels upon RBP expression could reflect changes in transcriptional or post-transcriptional processes since many of these RBPs function in RNA splicing. As expected, we did observe splicing changes in cells with acute RBP expression, but these changes were not associated with up-regulated genes (Figure S3), suggesting that altered splicing cannot account for the observed gene expression differences. These results show that, like cell-type specific TFs, RBP activators regulate endogenous transcription and may contribute to cell identity.

### RBP activators dynamically co-localize with transcriptional condensates

TFs can be recruited to genes by binding to preferred DNA sequence motifs or interacting with other protein factors^1,48–51^. By analogy, RBPs capable of regulating transcription could be localized to genes by interacting with RNA molecules synthesized from regulatory elements or the gene itself, or they could interact through protein-protein interactions with factors involved in transcription. To explore these possibilities, we generated dual-tagged murine embryonic stem cells (mESCs) where endogenous RBP activators were fused to the mStayGold fluorophore and Med1, a component of the Mediator coactivator complex and marker of transcriptional condensates^29^, was fused to the mScarlet3 fluorophore using CRISPR/Cas9 (Methods). To test whether the presence of RNA is important for RBP activator localization, mESCs were treated with the transcriptional elongation inhibitor DRB for 30 minutes followed by washout of the inhibitor. The RBP activator HNRNPH1 localizes diffusely throughout the nucleoplasm and to discrete puncta but relocated to nucleoli upon acute transcriptional inhibition (Figure 3A, Figure S4A). Proper nucleoplasmic localization was restored upon resumption of RNA synthesis during wash out of the inhibitor. These results suggest that HNRNPH1 localization depends on RNAPII-driven transcription of RNA molecules, which is one route enabling RBP localization to chromatin.

**Figure 3.**
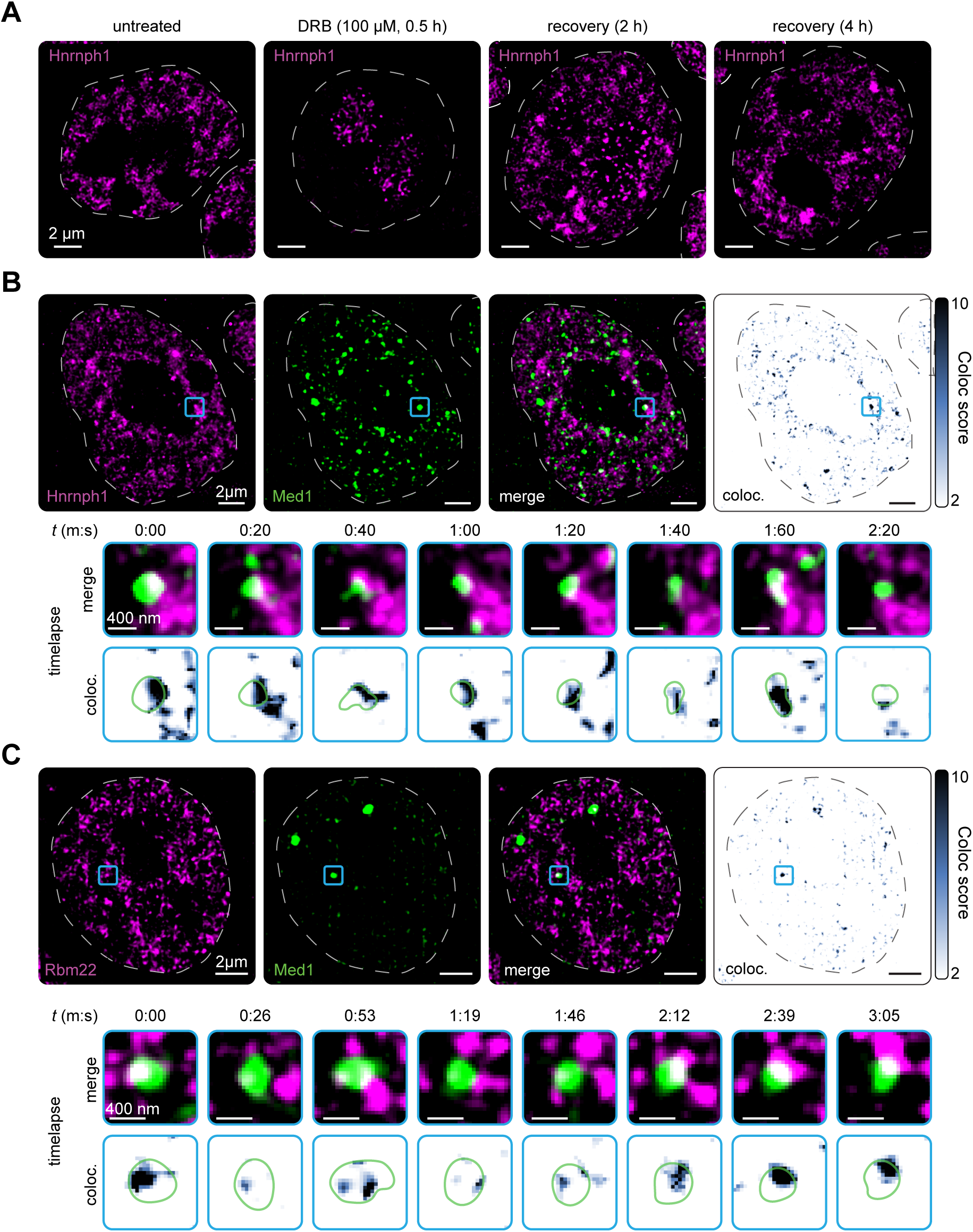
RBP activators dynamically co-localize with transcriptional condensates. A. Representative images exhibiting the change of Hnrnph1 nuclear localization upon DRB treatment and washout B. Top: Max intensity projection representative images and 3D colocalization heatmaps of cells with endogenous mScarlet-Med1 and mStayGold-Hnrnph1; Bottom: Representative zoom-in spots that shows the colocalized spots at different time frames. C. Top: Max intensity projection representative images and 3D colocalization heatmaps of cells with endogenous mScarlet-Med1 and mStayGold-Rbm22; Bottom: Representative zoom-in spots that shows the colocalized spots at different time frames.

To promote recruitment and loading of RNAPII, TFs can interact directly with the pre-initiation complex^52^ or recruit coactivators like the Mediator complex^53^. During transcription, progressive phosphorylation of the CTD of RPB1, the catalytic subunit of RNAPII, helps to recruit RBPs involved in RNA processing^54–57^, so we wondered if RBP activators could directly bind RNAPII. RBP activators were predicted to have more attractive interactions with RPB1-CTD than non-activators using force-field predictions of IDR-IDR interactions^58^ (Figure S4B). To test such interactions, co-immunoprecipitation of tagged RBP activators and blotting for RNAPII revealed that some factors, like HNRNPA1 and HNRNPH1, were able to bind RNAPII, whereas RBM22 was unable to do so, which was consistent with the force-field predictions (Figure S4B, S4C). These results suggest that some RBP activators may function through direct recruitment of RNAPII to active genes.

The dynamic interactions between TFs, RNAPII, coactivators, chromatin, and RNA molecules can promote the assembly of transcriptional condensates near genes^26,29,59–63^. RBPs and RNA molecules also form distinct condensates in the nucleus and cytoplasm, including nuclear speckles, paraspeckles, the nucleolus, stress granules, Cajal bodies, and others^64–66^. We compared RBP activation potential with force-field predictions of IDR-IDR interactions between RBPs and transcriptional coactivators, which revealed a switch in activation potential near the boundary of attractive and non-attractive IDR interactions (Figure S4D). We then sought to examine whether some RBP activator proteins could partition into transcriptional condensates, which would enable them to participate in transcriptional control. Re-examination of proteomics data^67^ revealed that many RBPs, including RBP activators, selectively partition into MED1 condensates formed in nuclear extract (Figure S4E). Live-cell imaging using lattice structured illumination super-resolution microscopy of dual-tagged mESCs showed that RBP activators Hnrnph1 and Rbm22 co-localized with transcriptional condensates marked by Med1 (Figure 3B, C). Timelapse imaging of these colocalized structures revealed that colocalization is dynamic, including transient partial mixing and interactions near the interface of transcriptional condensates and RBP-rich condensates (Figure 3B, C, Movie S1, Movie S2). These results show that RBP activators have the ability to partition into transcriptional condensates, but such interactions are dynamic and interfacial. It is possible that such interfacial enrichment reflects RNA-mediated de-mixing, which would be driven by differential affinities for RNA molecules between RBPs and transcriptional regulators^63,68–70^.

Together, the results are consistent with a model whereby RBP activators, like TFs, can be recruited to genes through both nucleic acid-mediated and protein-mediated interactions. The residence time of these interactions, though dynamic, are on the order of transcriptional events (seconds to minutes). Given these observations and predicted interaction potentials between RBPs and coactivators, we next sought to determine whether specific chemical features of RBP IDRs enabled their ability to activate transcription.

### RBP activators have distinct molecular grammars

TFs, coactivators, and RNAPII associate with each other through a combination of multivalent stereo-specific and chemoselective interactions. Specific avidities are driven by compatible molecular grammars, or the composition and linear patterning of amino acids within protein sequences^71^. For instance, TF activation domains are enriched with aromatic and other hydrophobic residues that are necessary for activation and thought to interact with hydrophobic regions in cofactors^72–74^. In other cases, alternating patterns of charged residues enable selective partitioning of transcriptional elongation factors in transcriptional condensates^67,75^. To determine if there are molecular features that distinguish RBP activators from non-activators, we compared their compositional and patterning enrichments relative to the human IDR-ome^76^. RBP activator IDRs were enriched in aromatic (F/Y) and polar (Q) residues, and in a subset of these factors, proline (P) residues (Figure 4A, Figure S5A). RBP activators were also depleted of basic residues (R/K). These molecular grammar enrichments and depletions were similar to what was observed for TF IDRs (Figure 4A, Figure S5B). In contrast, RBP non-activators were depleted of aromatic, polar, and proline residues and enriched for basic residues (Figure 4A). Enrichments and depletions were conserved among vertebrate orthologs (Figure 4B, Figure S6A). These enrichments and depletions, which correlate with activation activity (Figure S6B), suggest that RBP activators have a distinct molecular grammar compared to non-activators. The similarity of this grammar to TF activation domains may underlie the ability of RBP activators to stimulate transcription.

**Figure 4.**
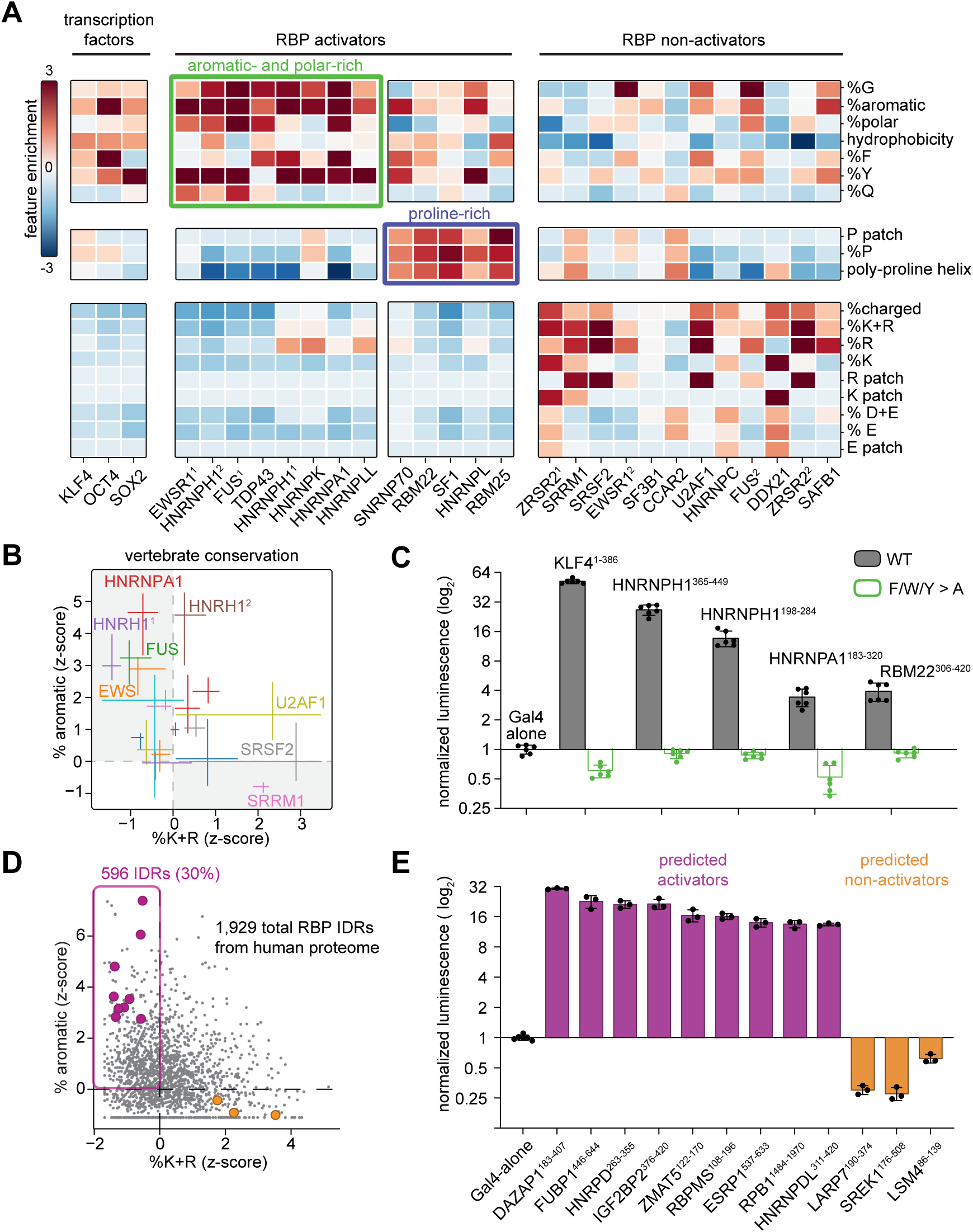
RBP activators have distinct molecular grammars. A. NARDINI+ analysis heatmap of amino acid composition of examples of transcription factors, RBP IDR activators and non-activators in Gal4 luciferase assay. B. NARDINI+ analysis of the enrichment of aromatic and basic residues of RBP orthologues across vertebrates. Error bars show the range of the top 95% similar orthologues. C. Bar plot depicting the normalized luminescence values for the Gal4 luciferase assay with WT or mutated IDRs (statistics in Table S1). D. Scatter plot showing the enrichment of aromatic and basic residues of all RBP IDRs with NARDINI+ analysis. Predicted and tested RBP IDR activators and non-activators in Figure 4E are colored. E. Bar plot depicting the normalized luminescence values for the Gal4 luciferase assay with predicted novel RBP IDR activators and non-activators based on the molecular grammar (statistics in Table S1).

We next sought to determine if enriched molecular grammars were important for RBP-mediated activation. Mutation of all aromatic residues to alanine in the IDRs of select RBP activators, which did not affect their expression level (Figure S1B), ablated their ability to activate a reporter gene in a manner similar to the TF KLF4 (Figure 4C, Figure S6C). Force-field analysis of IDR-IDR interactions predicted that loss of aromatic residues decreases attractive interactions between RBP IDRs and IDRs of the coactivators MED1 and BRD4 (Figure S6D), consistent with loss of reporter activation. These results show that grammar enrichments are necessary for RBP-mediated reporter activation.

Since our initial screen of RBPs only included factors with empirical evidence of chromatin enrichment, we reasoned that molecular grammar enrichments should be predictive of additional RBP activators encoded in the human proteome. Of the 1,929 predicted IDRs in RBPs across the human proteome^2^, 596 of them (30%) are enriched for aromatic residues and depleted of basic residues (Figure 4D, Table S3). We selected nine of these factors after filtering for factors that can localize to the nucleus, and each of these was able to significantly activate reporter expression in our Gal4-IDR assay (Figure 4E). In contrast, three selected factors that had grammars similar to our non-activators were unable to activate reporter expression (Figure 4E). These results show that molecular grammar enrichments are predictive of activation activity.

### RPB1-CTD activates transcription and recruits transcriptional coactivators

The prediction of additional RBP activators in the human proteome using molecular grammar enrichments included the CTD of RPB1 (Figure 4E). In humans, the CTD has 52 heptad (YSPTSPS) repeats, the first 26 of which are highly conserved and the last 26 more degenerate^77,78^ (Figure 5A). Molecular grammar enrichments of these repeats fit the aromatic-and proline-rich, charge-depleted grammar of RBP activators. Previous studies have found that CTD fused to Gal4 stimulates expression of a reporter gene^79,80^, and that this result was dependent on CTD length. We reproduced these efforts with our reporter system and found that full length CTD and the first 26 consensus heptads were able to stimulate reporter expression to a similar degree (Figure 5A, Figure S7A). However, truncation to 13 heptads or inclusion of the last 26 degenerate heptads abolished such activity. Mutating heptad tyrosines (Y) to alanine (A) similarly ablated activity, suggesting that aromatic residues are important for activation potential. Our results, together with previous efforts, are consistent with a non-enzymatic function for RNAPII, where the CTD is capable of stimulating transcription when recruited near a gene^53,77,80^.

**Figure 5.**
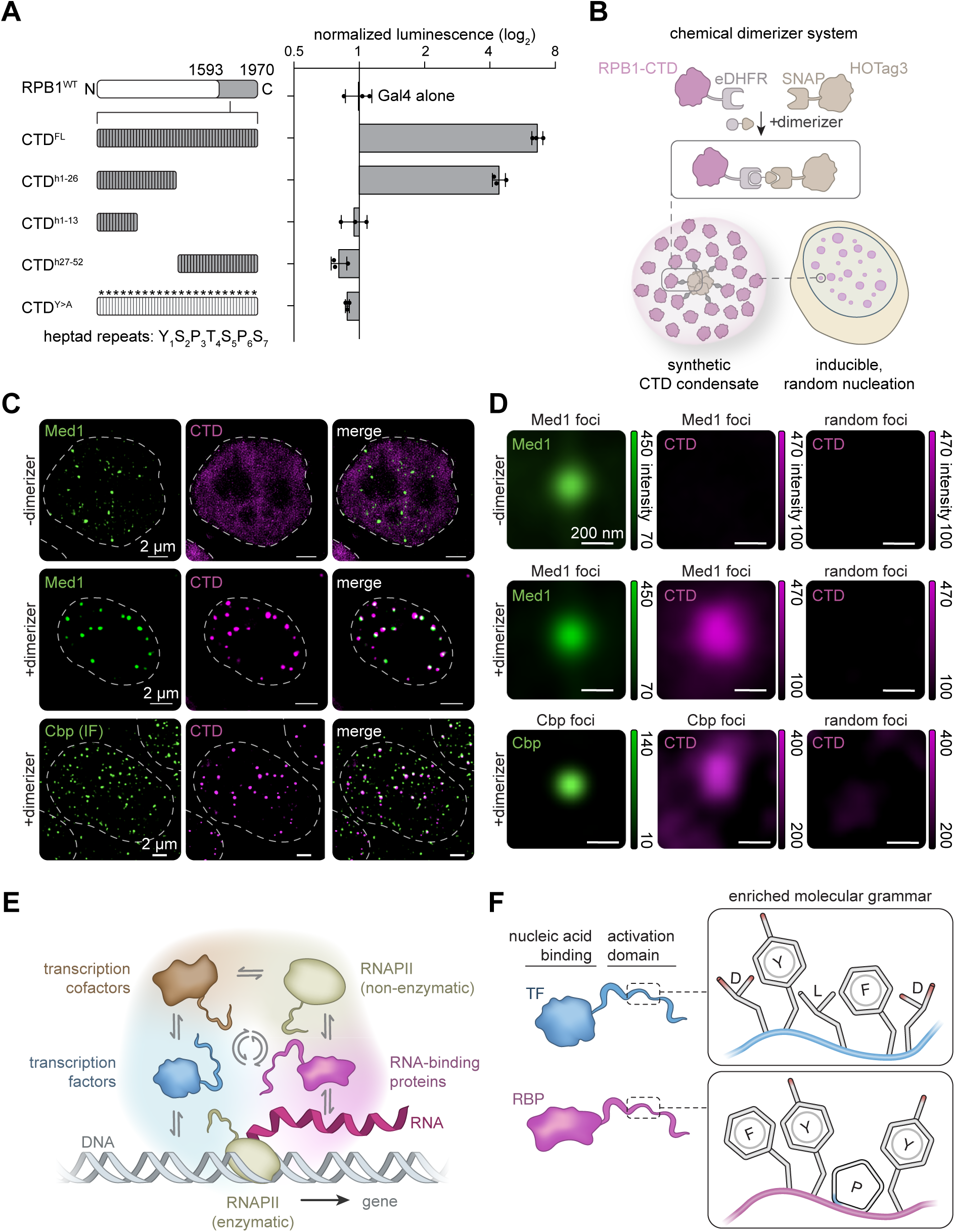
RPB1-CTD activates transcription recruits transcriptional coactivators. A. RPB1 CTD variants and corresponding Gal4 luciferase assay (statistics in Table S1). B. Schematic of the chemical dimerizer synthetic condensate system. C. Maximum projection of selected z slices of single nuclei with endogenous mScarlet-Med1 or antibody-stained CBP and CTD-mStayGold-eDHFR with or without dimerizer treatment. D. Radial distribution plots depicting the average intensity of CTD signal centered at Med1 or Cbp spots compared to control random nuclear areas. E. Depiction of protein-protein and protein-nucleic acid interactions that regulate transcription, including the enzymatic activity and non-enzymatic transcription activation ability of RNAPII. F. Depiction of molecular grammar enrichments in the activation domains of TFs and RBPs.

In mammalian cells, RNAPII complexes partition into transcriptional condensates and form clusters at active genes^62,81–83^. The number of RNAPII complexes at a particular locus can exceed the number of actively transcribing complexes^60^, and macro clusters of RNAPII do not always correlate with levels of nascent RNA^84^. Evidence that the CTD alone can activate transcription and the presence of RNAPII clusters near genes suggest that RNAPII itself could act as a transcriptional coactivator capable of recruiting transcriptional and chromatin regulators. To test this concept, we sought to generate CTD condensates in cells and measure their ability to partition transcriptional regulators. To avoid potential artifacts from simple over-expression systems, we used a chemical dimerizer system capable of forming condensates below bulk saturation concentrations in cells through recruitment of proteins to oligomeric seeds (Figure 5B)^85^. Cells expressing fusion proteins CTD-mStayGold-eDHFR and oligomeric scaffold protein SNAP-HOTag3-miRFP670 were treated with a small molecule (SNAP-TMP, Figure S7B) that induces dimerization. In the absence of dimerizer, the CTD-mStayGold-eDHFR signal remained largely diffuse throughout the nucleoplasm (Figure 5C, Figure S7C). With dimerizer treatment, the CTD-mStayGold-eDHFR fusion formed condensates (40 ±32 per cell, 208 ±45 nm mean diameter) that colocalized with SNAP-HOTag3 seeds, whereas mStayGold-eDHFR alone did not (Figure S7B). Induction of synthetic CTD condensates in Med1-mScarlet3 endogenously tagged cells revealed colocalization between synthetic CTD condensates and Med1 (Figure 5C, D). In a similar fashion, CTD synthetic condensates were also capable of recruiting the histone acetyltransferase and transcriptional coactivator CBP (Figure 5C, D). These results, together with transcriptional activation assays, are consistent with a model whereby RNAPII not only functions to catalyze RNA synthesis but can also act as a transcriptional coactivator capable of recruiting additional transcriptional regulators in a positive feedback loop (Figure 5E).

## Discussion

Here, we show that select RBPs commonly associated with post-transcriptional regulation harbor transcriptional activation domains that enable them to regulate transcription itself (Figure 5F). The RBP activation domains have defined molecular grammar enrichments that mediates this function. These enrichments are evolutionarily conserved, necessary for RBP-mediated gene activation, and are predictive of other RBP transcriptional regulators in the human proteome. Using molecular grammar enrichments, we found the CTD of RPB1, the largest subunit of RNAPII, can also recruit transcriptional coactivators to activate gene transcription, revealing a non-enzymatic feedback function for the RNAPII complex responsible for synthesizing all messenger RNAs. Together, these results reveal a non-canonical role for RBPs in transcriptional control and position select RBPs alongside established TFs and cofactors.

Accumulating evidence suggests that RBPs play dual roles in regulating both RNA synthesis and co-transcriptional RNA processing^6,86–91^. These biochemical functions are closely linked and likely coupled, as most RNA splicing events occur within proximity (<300 nm) of their encoding gene^92,93^. Genes that are transcribed near nuclear speckles, bodies enriched with RNA-binding proteins, are more efficiently spliced^34^ and transcribed^94^. Enhanced splicing is due to increased local concentrations of splicing factors, but how proximity to nuclear speckles or the presence of introns stimulates nascent transcription has largely remained a mystery^95,96^. Our results suggest that some RBPs, particularly regulators of RNA splicing, can locally enrich at gene regulatory elements to stimulate transcription in a fashion similar to TFs and cofactors.

Similar to TFs, RBPs localize to genes through binding to nucleic acid and other transcriptional regulators. Live imaging of RBP activators and transcriptional components revealed dynamic colocalization between these proteins, and such colocalization enriched near the interface of transcriptional and RBP condensates. Differential affinities between protein-protein and protein-RNA interactions can create interfacial, multiphasic condensates governed by local RNA concentration in simplified, equilibrium-driven systems^69^. Since RBP activators harbor canonical RNA-binding domains as well as IDRs capable of interacting with transcriptional proteins, the observed interfacial organization could be understood in terms of classical coacervation theory, where rising concentrations of RNA near an active gene during transcription lead to partial condensate mixing at the interface. These observations are consistent with recent findings that partitioning of the RBP PSPC1 can promote RNAPII transcription^97^. The observed interfacial organization can also be understood by analogy to the many nuclear condensates that exhibit core and shell architectures, including nuclear speckles^98^, paraspeckles^99^, histone locus bodies^100^, and the nucleolus^101^. The nucleolus is particularly relevant, as factors involved in pre-ribosomal RNA transcription and pre-ribosomal RNA processing form RNA-dependent, multiphasic networks composed of nested, interfacial condensates^102,103^. The far greater number of DNA- and protein-associated components in the nucleolus likely underlies why its fibrillar and dense fibrillar layers exhibit more stable organization than the highly dynamic spatiotemporal patterns we observe for RNAPII-associated condensates.

Our results show that RBPs activate gene transcription through molecular grammar enrichments in their IDRs that partially mirror TF activation domains. Recent studies have demonstrated that compatible molecular grammars enable multivalent interactions that are important for specific enrichment of transcriptional components^67,104^. Through force-field predictions and experiment, we find that enrichment of aromatic, polar, and proline residues and depletion of basic residues contribute to RBP activation potential and interaction with transcriptional coactivators. High throughput efforts to identify molecular grammar enrichments for TF activation domains have found that these domains tend to be enriched with hydrophobic, aromatic, and acidic residues^41,105^. The conventional model is that acidic residues help expose adjacent hydrophobic residues for interactions with cofactors^73^. We did not observe enrichment of acidic residues in RBP cofactor IDRs, but the enrichments we did observe support a model whereby RBP activators use pi-stacking and cation-pi interactions to recruit transcriptional cofactors. Molecular grammar enrichments were predictive of additional activators in the human proteome, with estimates suggesting there could be hundreds of putative RBP activators (Table S3) complementing the ∼1600 human TFs and cofactors encoded in the human genome^1^. The complexity and diversity of mechanisms governing RBP-mediated transcriptional regulation will likely match diverse ways by which TFs and cofactors specifically regulate transcription initiation and elongation.

Molecular grammar enrichments led us to consider the CTD of RPB1, which has been comprehensively studied as a regulator of RNA synthesis and processing^106^. Canonical models of gene regulation typically consider a spatially constrained view of CTD-mediated function, where the RNAPII holoenzyme is tethered to the genome during RNA synthesis. Early studies found that fusion of the CTD to heterologous DNA-binding domains could activate transcriptional reporters, and this effect was dependent on CTD length^79,80^. Tethering to DNA was thought to stimulate CTD phosphorylation, which is a critical regulatory toggle for transcription initiation, elongation, and assembly of the RNA spliceosome^57,107^. Our results, combined with observations that active genes can harbor RNAPII complexes not engaged in transcription^60,84,108^, support an additional, non-enzymatic function for RNAPII in a regulatory feedback loop. Synthetic nuclear condensates scaffolded by CTD were capable of recruiting multiple transcriptional cofactors. The molecular grammar of the CTD is important for this function, as mutation of critical tyrosine residues abolishes activation potential and prevents proper condensate formation^109^. In this model, RNAPII complexes that partition into transcriptional condensates could recruit additional cofactors through compatible molecular grammar interactions with the CTD, which would reinforce or stabilize gene-specific transcription.

Evidence that RBPs regulate transcription has implications for both fundamental models of gene control and gene dysregulation in disease. Enhancer and promoter elements are transcribed to RNA molecules, some of which have been implicated in transcriptional regulation^110–113^. Our results suggest that this could be in part through early recruitment of RBP activators. Active genic transcription creates high RNA concentrations at gene loci, where nascent transcripts are bound by multiple protein factors that form spatial compartments contributing to genome structure and gene expression^33,114^. Our findings raise the possibility that RBP-RNA networks analogous to TF-DNA networks exist as part of a cis regulatory code for transcriptional control. RBPs as transcriptional activators could also inspire alternative mechanisms in the understanding of diseases driven by RBP mutations. For instance, core spliceosomal proteins are recurrently mutated in hematological disorders, and RNA sequencing studies have found disease-associated alterations in both transcription and RNA splicing^20^. Mutations in some RBP activators, including FUS, EWSR1, and TDP43, are frequently associated with neurodegenerative disorders, where pathological aggregates accumulate in the cytoplasm. Our results suggest that such cytoplasmic sequestration could contribute to disease pathogenesis through loss of proper transcriptional control, in line with current models^115,116^. Collectively, these findings reposition select RBPs and RPB1 as transcriptional regulators in addition to their post-transcriptional roles, expanding the molecular logic that governs gene expression.

### Limitations of the Study

Since our reporter assays used DNA-based recruitment of RBP IDRs to probe their function in transcription, it is important to note that we cannot exclude the possibility that some non-activator RBPs nevertheless regulate transcription through alternative mechanisms. For instance, the IDRs of splicing regulators SRSF2 and SF3B1 did not activate reporter expression, but prior evidence supports their ability to regulate nascent transcription, perhaps through promotion of elongation^18–20^. Moreover, as is the case for TFs, it can be difficult to parse direct transcriptional regulation from steady-state, population-averaged RNA sequencing data. It’s possible that some of the differences in differential expression observed in our studies can be attributed to differences in transcription, post-transcriptional processing or degradation, or both processes simultaneously. We attempted to mitigate possible secondary, non-transcriptional effects through acute measurement and analysis of post-transcriptional effects.

## Methods

### Data and code availability

RNA sequencing data has been deposited to GEO (GSE327996). Other data and scripts used for analysis are available upon reasonable request.

### Cell culture

HEK293T cells were cultured at 37°C with 5% CO2 in a humidified incubator on tissue-culture plates containing culture made according to the following recipe: 450 mL DMEM medium (ThermoFisher Cat. 11965118), 50 mL FBS (ThermoFisher Cat. 16140071), and 5 mL penicillin-streptomycin (Life Technologies, 15140163; stock 10^4 U/mL). Cells were passaged by TrypLE (Life Technologies, 12604021) incubation for 3 minutes. The digestion reaction was quenched by adding the culturing medium made by the recipe above. Cells were passaged every 2-3 days.

K562 cells were cultured at 37°C with 5% CO2 in a humidified incubator in suspension flasks containing culture medium made according to the following recipe: 450 mL RPMI-1640 medium (ThermoFisher Cat. 11875119), 50 mL FBS (ThermoFisher Cat. 16140071), and 5 mL penicillin-streptomycin (Life Technologies, 15140163; stock 10^4 U/mL).

mESCs were cultured at 37°C with 5% CO2 in a humidified incubator on 0.2% gelatinized (Sigma, G1890) tissue-culture plates in the medium made according to the following recipe: 960 mL DMEM/F12 (Life Technologies, 11320082), 5 mL N2 supplement (Life Technologies, 17502048; stock 100X), 10 mL B27 supplement (Life Technologies, 17504044; stock 50X), 5 mL L-glutamine (GIBCO 25030-081; stock 200 mM), 10 mL MEM nonessential amino acids (GIBCO 11140076; stock 100X), 10 mL penicillin-streptomycin (Life Technologies, 15140163; stock 10^4 U/mL), 333 μL BSA fraction V (GIBCO 15260037; stock 7.50%), 7 μL β-mercaptoethanol (Sigma M6250; stock 14.3 M), 100 μL LIF (Chemico, ESG1107; stock 10^7 U/mL), 100 μL PD0325901 (Stemgent, 04-0006-10; stock 10 mM), and 300 μL CHIR99021 (Stemgent, 04-0004-10; stock 10 mM). To passage the cells, 1X PBS (Life Technologies, AM9625) was used to wash the cells, followed by TrypLE (Life Technologies, 12604021) treatment for 3 minutes. The digestion reaction was quenched by adding serum-containing media made by the following recipe: 425 mL DMEM KO (GIBCO 10829-018), 5 mL MEM nonessential amino acids (GIBCO 11140076; stock 100X), 5 mL penicillin-streptomycin (Life Technologies, 15140163; stock 10^4 U/mL), 5 mL L-glutamine (GIBCO 25030-081; stock 100X), 4 μL β-mercaptoethanol (Sigma M6250; stock 14.3 M), 50 μL LIF (Chemico, ESG1107; stock 10^7 U/mL), and 75 mL of fetal bovine serum (Sigma, F4135). Cells were passaged every 2 days.

### Inducible, acute expression of Halo-tagged RBPs in K562 cells

Piggybac transposon-compatible plasmids^10^ were generated by cloning RBP gene sequences (PCR from K562 cDNA) into doxycycline-inducible, N-terminal Halo-tagged expression construct with a hygromycin-resistance gene using NEBuilder HiFi DNA Assembly (NEB, E5520S). For transfection, 250K wildtype K562 were plated in 6-well plates and simultaneously transfected with 1 μg of the expression vector and 1 μg of the PiggyBac transposase (purchased from System Biosciences) using Lipofectamine 3000 (ThermoFisher, L3000075), according to manufacturer instructions. The next day, medium was replaced by medium supplemented with 500 μg/mL hygromycin (ThermoFisher, 10687010) for selection. Un-transfected control cells were used to assess selection.

#### RNA-seq analysis

For RNA-seq, expression of RBPs was induced for 4 hours using 100 ng/mL doxycycline (Sigma, D9891-5G) for all factors except for RBM22 (10 ng/mL) in biological duplicates. Doxycycline doses were determined by titration and measurement of RBP expression by qRT-PCR. As a control, we included a K562 cell line with stable expression of the Halo tag alone treated with doxycycline to control for any effects due to doxycycline treatment and induction of the integrated expression vectors. RNA-seq quality control and sequencing was carried out by Novogene (20-30 million paired-end reads per sample, 150nt). The quality of raw fastq files were checked with *fastqc* [https://www.bioinformatics.babraham.ac.uk/projects/fastqc/]. After trimming the adaptors with cutadapt^117^, pair-end reads were aligned to the reference genome using *STAR*^118^ with the ENCODE options in *STAR* manual 2.7.0a by Dobin in 2019. *samtools*^119^ was used to sort and index the bam files using default parameters, and the count table was generated by running *featureCounts* using default parameters^120^. Differential gene expression analysis was performed with *DESeq2* using default parameters in RStudio^121^. In Figure 2A, z score heatmap was generated using *pheatmap* in RStudio. In Figure S2A-S2E, volcano plots were generated using *EnhancedVolcano* in RStudio. In Figure 2B and Figure S2F, upset plots were generated using *upsetplot* in python In Figure S3, splicing analysis was performed by *rMATS turbo* using default parameters^122^. Volcano plots were generated using *matplotlib* in python.

#### ChIP-seq analysis

For Figure S1A, fold-change bigWig files from ChIP-seq experiments targeting activator RBPs were obtained from the ENCODE portal^123^ (https://www.encodeproject.org/). Accession numbers for the files used for heatmap analysis were as follows: ENCFF251SRH, ENCFF313ZXN, ENCFF879COR, ENCFF639KRR, ENCFF209WQK, ENCFF537MTV, ENCFF338KQK, ENCFF281TJW, ENCFF937ZFO, ENCFF727KHF, ENCFF499TQE. Enhancer and super-enhancer locations in K562 cells from Hnisz et al. were used for downstream analysis^47^. The *deeptools* package^124^ was then used to generate heatmaps showing ChIP-seq enrichment at each enhancer and super-enhancer with 1 kB padding. Deeptools computeMatrix was run with parameters scale-regions and plotHeatmap was run with --zMin 0 --zMax 4.0, with all other parameters remaining at their default values. For the promoter heatmaps, the *deeptools reference-point* command was used with 500 bp padding. Transcription start site locations were obtained from refSeq for all hg38 gene isoforms. The *plotHeatmap* function was used with the same parameters as before on the output of the reference-point.

For Figure 2C, ChIP-seq data was used to validate RBP localization at super-enhancer-associated genes in RBP overexpression experiments. Upregulated and neutral genes from the OE experiment were intersected with a list of genes associated with super-enhancers in K562 cells^47^. Profile plots centered around upregulated and neutral super-enhancer-associated genes using RBM22, FUS, and RBM25 ChIP bigWig files were created using the *deeptools package*. Deeptools computeMatrix was run with 5000 bp padding, scale-regions, and --regionBodyLength 5000. The plotProfile command was then run with –sortRegions keep and – outFileNameData to extract the underlying data for the plot. All other parameters for both commands were kept as default. To smooth out the graph, a moving average of every 5 data points within the profile plot generated data was calculated and used to regenerate the plots. Upregulated genes were defined as log2fc > 1 in the OE experiment, while neutral genes were defined as ™0.5 < log2fc < 0.5.

### Prediction of intrinsically disordered regions of RBPs

For Figure 1B, Figure 4A, B, and D, Figure S5A, Extended Figure 6A-C the intrinsically disordered regions (IDRs) of select chromatin-enriched RNA binding proteins were predicted by *metapredict* (version: 3.0.1), using their default parameters for identifying IDRs OR using a threshold of 0.5 predicted disorder and filtering on contiguous regions^125^.

### Gal4 luciferase transcription reporter assay

Predicted IDR sequences (see “**Prediction of intrinsically disordered regions of RBPs**”) were amplified using PCR from plasmids (Addgene: #65926, #17578, #10820, #99021, #141327, #148120, #46958, #64925, #128803, #35096, #64923, #82576, #23026), or cDNA generated from RNA isolated from wildtype K562 cells (Zymo Quick-RNA Miniprep Kit (R1055) for RNA isolation and Superscript IV (Thermo Scientific, 18090200) with oligodT primers (Thermo Scientific, 18418020) for cDNA generation). RBP IDRs were cloned into an empty vector fusing them to a Gal4 DNA-binding domain downstream of the hUbiC promoter and a kozak sequence using Gibson cloning strategy following the manufacturer’s instructions (NEB, E5520S). A luciferase reporter construct^26^ containing firefly luciferase downstream of six UAS repeats followed by a minimal CMV promoter, and a renilla transfection control (pRL-SV40, Promega) were used. For Figure 1D, Figure 4C and E, Figure 5A, 200k per well of HEK293T cells were plated in 24-well plates the day before transfection. 150ng Gal4-IDR, 100ng firefly luciferase, and 50ng renilla luciferase plasmids were transfected into HEK293T cells in triplicate using Lipofectamine 3000 Transfection following kit instructions (Thermo Scientific, L3000150). “No Gal4 condition” refers to the transfection of a plasmid that expresses the bacterial transcription factor LacI which is not able to bind the UAS sequences to the cells. After 4 hours of incubation, cells were lysed and luminescence was measured in white bottom 96-well plates (Corning, 07-200-589) using the Dual-Luciferase Reporter Assay System (Promega, E1960) following the manufacturer’s instructions.The assays were conducted on 3 biological replicates per genotype, and luminescence readings were measured in technical duplicates (BioTek Synergy H1 Multimode Reader). The firefly luminescence values of each condition were first normalized to the renilla luminescence and then compared to the Gal4-alone control using one-Way ANOVA (Table S1).

For mutant versions of RBPs in Figure 4E, all aromatic residues (W/F/Y) or all proline residues (P) were mutated to alanine. Plasmids were constructed using Gibson cloning (NEB, E5520S) with gBlocks (IDT) encoding the mutations with codon optimization. For CTD truncations in Figure 5A, plasmids were constructed using Gibson cloning with PCR fragments using specific primers that amplified truncations of the CTD. For the Y>A CTD mutation, a plasmid was constructed by Gibson cloning using a gBlock where each Y in each heptad was mutated to alanine and codon-optimized. Transfection and luciferase assay were carried out the same way as in Figure 1D.

For the Gal4 luciferase assay in Figure S1C, the luciferase reporter plasmid with a strong CMV promoter was used to examine the repression ability of RBP IDR non-activators. Transfection and luciferase assay were conducted following the same protocol as Figure 1D.

For RT-qPCR in Figure S1D-S1E and Figure S7A, transfection was carried out the same way as the luciferase assay. After 4h incubation, RNA was extracted using ZYMO Quick-RNA Miniprep Kit (R1055). RT-qPCR was performed with PowerTrack™ SYBR Green Master Mix (A46109) with primers: DMF076_qPCR_hsBACT_F, DMF077_qPCR_hsBACT_R, DMF093_qPCR-GAL4-F1, DMF094_qPCR-GAL4-R1, either DMF085_qPCR-FLUC-F1 and DMF086_qPCR-FLUC-R1 (Figure S1D and S1E), or DMF091_qPCR-FLUC-F4 and DMF092_qPCR-FLUC-R4 (Figure S1D and S1E, Figure S7A, Table S4). The assays were conducted on 3 biological replicates per genotype. The firefly luciferase Ct values of each condition were first normalized to the Ct value of beta-actin, and then compared to the Gal4-alone control using one-way ANOVA(Table S1)

### Endogenously tagged cell line generation

CRISPR/Cas9 was used to generate mESC RBP and Med1 endogenously tagged cell lines. RBP was tagged with FKBP (F36V)-HA-mStayGold, and Med1 was tagged with HA-mScarlet3. sgRNAs targeting the N-terminus of proteins were cloned into the vector expression Cas9 and puromycin (NEB, M0202M) (Table S4). The sgRNA sequences we used in the experiments are: Hnrnph1 - 5’ TCCCAGCATCATCGTCCCTT 3’; Rbm22 - 5’ ACGCGGTGAGTGCACGCCGC 3’; Med1 - 5’ TGTCAGGATGAAGGCTCAGG 3’ ^29^. Repair templates, including a 5’ homologous arms, fluorescent tags, and 3’ homologous arms, were cloned into pUC19 vector using Gibson cloning (NEB, E5520S). To generate dual tagged cell lines simultaneously where either Hnrnph1 or Rbm22 is tagged with FKBP (F36V)-HA-mStayGold and Med1 is tagged with HA-mScarlet3, 300k mESCs were transfected with sgRNA and repair template plasmids for both RBP and Med1 (600ng each) using lipofectamine. Two days after transfection, cells were selected with puromycin (1:5000 dilution; Gibco, A1113803) for 2 days. One week later, cells were sorted based on mStayGold and mScarlet3 fluorescence.

### Super-resolution microscopy and transcription inhibition

For live cell images in Figure 3, Figure 5, and Figure S7B, 35mm glass bottom imaging dishes (Mattek, Part No: P35G-1.5-20-C) were coated with poly-L-ornithine (Sigma, P4957-50ML) and laminin (Coring, CB-40232) and incubated in the humidified incubator overnight. 250k mESCs were seeded to the plate the next day, and images were taken 1-2 days after. For fixed cell imaging in Figure 5C (CBP), cells were seeded first on glass coverslips before fixation. Cells were imaged with the Zeiss Lattice SIM 5 super-resolution microscope system (camera: Dual Orca Fusion BT; objective: Plan-Apochromat 63x/1.40 Oil DIC M27; illumination wavelength: 561 for Med1, 488 for RBPs; number of SIM phases:13 for z stack imaging, and 9 for timelapse imaging; z-step: 0.12um or 0.36um in Zeiss “Leap” mode; exposure time: 20ms; laser power: 10%-30%; processing method: SIM^2^ with default parameters). To correct for dual camera alignment, fluorescent beads (Invitrogen, T7280) were used to align the cameras, and images of the beads were taken with the same imaging conditions as ground truth for channel alignment via Zen Black Software. Timelapse imaging was bleachcorrected in ImageJ.

For transcription inhibition in Figure 3A and Figure S4A, cells were treated with 100uM DRB (SigmaAldrich, D1916-10MG for 30min, and imaged immediately after that. For the washout experiment, cells were first treated with DRB for 30min. DRB was then washed out by PBS and the medium was replaced with 2i++. Cells were imaged 1h, 2h, and 4h after DRB washout.

Max projection of selected contiguous z stacks was used to generate the representative images in Figure 3 to capture the whole volume of the zoom-in condensates.

### Colocalization heatmap

Representative timelapse z-stack images were selected for each dataset to generate colocalization heatmaps. The intensity of each channel was first z-score standardized by subtracting the mean and dividing by the standard deviation of the image. We defined a colocalization metric to highlight colocalized pixels between both channels while mitigating spurious colocalization arising from extreme values in one channel:

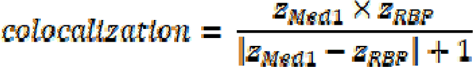

Colocalization heatmaps were generated using the Python package *matplotlib*. All heatmaps are displayed on the same scale.

### Radial distribution plot

Images were first converted to maximum intensity projections along the z-axis. Condensates were then segmented using a Laplacian of Gaussian transform with a symmetric σ of 2 and a minimum threshold of 8 standard deviations above the mean. Then nuclei were segmented using Cellpose-SAM^126^ with the following parameters: diameter 300 pixels, flow threshold 0.4, and cell probability threshold 0.0. Condensates outside these regions were discarded. Then 1μm crops were collected, centered on the centroid of each condensate of a particular channel. Fluorescence intensities were then averaged across all crops for each pixel within the 1μm box, showing the average spatial distribution of intensities of one channel compared to the other.

### Molecular grammar analysis of intrinsically disordered regions

For Figure 4A, B and D, Figure S5, Figure S6A and S6B, To quantify the relative enrichment of the molecular grammar of different RBP IDRs, the Google Colab notebook were used: https://github.com/kierstenruff/RUFF_KING_Grammars_of_IDRs_using_NARDINI-?tab=readme-ov-file. Briefly, the IDR sequences we tested in the Gal4 luciferase assay or transcription activation domains from a previous highthroughput assay^27^ were stored as a fasta file, which was used as the input to run the NARDINI+ analysis. The output result files contain the raw value and the z-score of different amino acid features of each IDR sequence. The z-score file was used to generate the heatmaps showing the molecular grammar enriched in TF activators and RBP activators.

### Synthetic condensate cell line generation

Using NEBuilder HiFi DNA Assembly (NEB, E5520S), a protein or IDR of interest was cloned into a doxycycline-inducible, Piggybac transposon-compatible plasmid with hygromycin resistance and eDHFR in the backbone. A separate doxycycline-inducible, Piggybac transposon-compatible plasmid was cloned to include neomycin resistance, HOTag3, and 3XSnapf. For transfection, 500k cells of either WT mESCs of sorted mScarlet3-MED1 mESCs were plated per well of a 6-well plate. Invitrogen Lipofectamine 3000 Transfection Reagent (Thermofisher Scientific, L3000075) was used to transfect 1000ng of DNA of each of these plasmids, as well as 1000ng of a Piggybac transposase plasmid per well at the same time cells were seeded in Serum/LIF media. One well was left untransfected to serve as a control during selection. After 4 hours at 37°C incubation, the Serum/LIF media was replaced with 2i++ media, and the cells were left to recover overnight. The next morning, the cells were split 1:1 into 10cm plates in 1:100 hygromycin (Thermofisher Scientific, 10687010) and 1:250 geneticin (Thermofisher Scientific, 10131035) in 2i++ media. Every day, a 1X PBS wash was done on the cells, and the media was replaced with fresh selection media until all control cells were dead, at which point the newly generated cell lines were PBS-washed and placed in regular 2i++ media to expand.

### Synthetic condensate formation

For Figure 5C, doxycycline-inducible wild type or sorted mScarlet3-Med1 mESC cells endogenously expressing both HOTag3 fused to 3XSnapf and eDHFR fused to a protein or IDR of interest were seeded 100k cells per dish in 35mm glass-bottom imaging dishes coated with 1:20 poly-L-ornithine in 1X PBS and 1:20 laminin in 1X PBS. Cells were incubated at 37°C for 2 days until small colonies formed. 1µg/mL dox was added to each dish and incubated for 2 hours. The dimer snap/TMP was added to a working concentration of 1µM per dish; alternatively, the same volume of DMSO (Fisher BioReagents, BP231100) was added to a negative control dish. Cells were incubated 2 more hours before aspirating the media from each dish, washing with 1X PBS, and replacing with phenol red free 2i- - media. Cells were then imaged using Zeiss Lattice SIM 5 super-resolution microscope.

For colocalization of synthetic CTD condensates and CBP in mESCs in Figure 5, 20k mESCs were seeded on coverslips coated with 1:20 poly-L-ornithine in 1X PBS and 1:20 laminin in 1X PBS in a 24-well plate. Cells were incubated at 37°C for 2 days before treating with dox and the dimer snap/TMP, as described above, and fixing with 4% PFA (avantor, BT140770-10X10). Cells were then permeabilized in 0.5% Triton X100 (Sigma, EM-9410) in PBS (PBST) for 10 minutes, and blocked with 4% IgG-free Bovine Serum Albumin (avantor, VMR 102643-518) in PBST for 1 hour. Washes were performed between each step with PBS. CBP/KAT3A/CREBBP antibody (Santa Cruz sc-7300) was diluted 1:50 in 4% BSA, and cells were incubated overnight in a humidity chamber at 4°C. The following day, cells were washed with PBS and Donkey anti-Mouse IgG (H+L) Highly Cross-Adsorbed Secondary Antibody with Alexa Fluor 555 (Invitrogen, A31570) diluted 1:500 in 4% BSA, was added. The cells were incubated for 1 hour, followed by three washes in 1X PBS. Cells were mounted on slides using vectashield (VWR, 101098-042) and imaged using Zeiss Lattice SIM 5 super-resolution microscope.

### IDR interactivity analysis

For Figure S4B and S4D, and Figure S6D, FINCHES^58^ was used with the Mpipi forcefield to calculate the epsilon scores between RBP IDRs tested in the GAL4 luciferase assay and cofactors MED1 and BRD4. Epsilon scores represent the average interactivity of the two IDRs, with negative values indicating attraction, positive indicating repulsion. Graphs were plotted with the negative epsilon scores and labeled as either activators or nonactivators based on the GAL4 luciferase assay. Mean epsilon scores were also predicted for interactions between RBPs and RPB1-CTD, which were compared between activators and non-activators using box plots.

### Molecular grammar conservation

Vertebrate orthologues (ID: 7742) of RBPs were retrieved from OrthoDB^127^ by Uniprot ID. To map conserved residues, orthologues were first aligned to the human sequence using multisequence alignment by Clustal Omega^128^ with default parameters. We then used NARDINI+ (refer to “Molecular grammar analysis of intrinsically disordered regions”) to calculate z-score enrichments for all orthologues and plotted the 95% confidence interval of z-score enrichments in Figure 4B and Figure 6A.

### Western blot

For Figure S1B, 200k per well of HEK293T cells were plated in 24-well plates the day before transfection. 1000ng Gal4-IDR plasmids (Gal4-IDR protein has an HA tag) were transfected into HEK293T cells using Lipofectamine 3000 (ThermoFisher Cat. L3000150). 24h after transfection, cells were lysed with RIPA Lysis and Extraction Buffer (Thermo Scientific, 89901) and Halt™ Protease and Phosphatase Inhibitor Cocktail (Thermo Scientific, 78440) on ice for 25min. The lysate was sonicated for 10s and treated with DNase I(ZYMO, E1010) for 15min at RT. The lysate was then spinned down and supernatant was taken for measuring protein concentration using Pierce™ BCA Protein Assay Kit (Thermo Scientific, 23250). After standardizing the protein concentration, DTT (final concentration 0.05M, BIO-RAD, 1610611) and Pierce™ Lane Marker Non-reducing Sample Buffer (5X) (VWR, PIER39001) were added to the lysate. The lysate was boiled for 5min at 95°C to denature the proteins and placed back on ice. Lysate was then loaded to a 4–15% Mini-PROTEAN® TGX™ Precast Protein Gel (BIO-RAD, 4561086). The gel was run for 30-40min at 150V in 1x TGS running buffer (diluted from 10x Tris/Glycine/SDS (BIO-RAD, 1610732) in diH2O). Before transferring the proteins, the filter papers (BIO-RAD, 12023835) were immersed in the Trans-Blot transfer buffer (5X TurboBlot transfer buffer (BIO-RAD, 10026938) and 20% methanol (final concentration) in diH2O). PVDF membranes (BIO-RAD, 12023927) were equilibrated in 100% methanol for 2-3min and rinsed with the Trans-Blot transfer buffer. Protein transfer was carried out with the Trans-Blot Turbo Transfer System (BIO-RAD, 1704150) for 7min at 25V. The membranes were washed with 1x TBST (20x TBST (Thermo Scientific Chemicals, J77500.K2) diluted in diH2O) for 5min, twice. The membranes were then blocked with EveryBlot Blocking Buffer (BIO-RAD, 12010020) at RT for 30min. The membranes were then incubated with primary antibodies (1:10000 HRP anti-beta actin antibody, abcam, ab20272; HA tag Polyclonal antibody, Proteintech, 51064-2-AP) overnight at RT. Membranes incubated with the HA tag Polyclonal antibody were immersed in 1:10000 Goat anti-Rabbit IgG (H+L) Secondary Antibody, HRP (Invitrogen, 31460) for 30min. All membranes were washed with 1xTBST for 5 times, 5min each. SuperSignal™ West Femto Maximum Sensitivity Substrate (Thermo Scientific, 34095) was used for chemiluminescent detection. Membranes were imaged using ChemiDoc MP Imaging System (BIO-RAD).

### Co-immunoprecipitation

#### Cell Culture and Doxycycline Induction

K562 cell lines containing doxycycline-inducible constructs for HNRNPH1, SRSF2, HNRNPK, HNRNPA1, SNRNP70, and RBM22 were maintained in RPMI 1640 medium supplemented with Fetal Bovine Serum (FBS) and penicillin/streptomycin. Cells were cultured at 37°C in a humidified 5% CO2 incubator and passed three times post-thaw to reach optimal density prior to induction. For induction, cells were seeded at a density of 1 × 10L cells/mL. Cultures were treated with either doxycycline (Dox+) using a 1000X stock solution or an equivalent volume of ultrapure water for the uninduced controls (Dox-). Cells were incubated for 6 hours prior to harvesting.

#### Cell Harvest and Protein Extraction

Post-induction, cells were pelleted by centrifugation at 1000 × g for 5 minutes and the media was aspirated. Cell pellets were washed once with 1 mL of cold PBS, centrifuged again at 1000 × g for 5 minutes at 4°C, and thoroughly drained. Pellets were resuspended in 500 µL of Co-IP Lysis Buffer (Pierce, 88804) supplemented with phosphatase and protease inhibitors (PPi) (Thermo Scientific, 78440). The lysates were incubated on ice for 5 minutes with periodic mixing, followed by centrifugation at 14,000 × g for 10 minutes to clear cell debris. The resulting supernatant was collected. Total protein concentration was determined using a Pierce BCA Protein Assay Kit (Thermo Scientific, 23250), with samples and standards incubated at 37°C for 30 minutes before reading absorbance at 562 nm on a microplate reader (BioTek Synergy H1 Multimode Reader).

#### Co-Immunoprecipitation (Co-IP)

Co-immunoprecipitation was performed as described in the manufacturers protocol for Pierce™ Classic Magnetic IP/Co-IP Kit (cat.88804). Briefly, immune complexes were prepared by combining 500–1000 µg of total protein lysate with 2–10 µg of anti-HA antibody (Abcam ab9110). The volume was adjusted to 500 µL using IP Lysis/Wash Buffer (Pierce, 88804). The mixture was incubated with gentle agitation for 1–2 hours at room temperature, or overnight at 4°C. Pierce Protein A/G Magnetic Beads (0.25 mg, 25 µL per sample, Pierce, 88803) were pre-washed with IP Lysis/Wash Buffer and collected using a magnetic stand. The pre-washed beads were added to the immune complex and incubated for 1 hour at room temperature with rotation. Following incubation, the unbound supernatant was removed and saved for analysis. The beads were washed three times with 500 µL of IP Lysis/Wash Buffer (Pierce, 88804) and once with 500 µL of ultrapure water. Target proteins were eluted by incubating the beads with 100 µL of Low-pH Elution Buffer for 10 minutes at room temperature. The eluate was magnetically separated and immediately neutralized with 10 µL of Neutralization Buffer. Cell lysates were transferred to fresh tubes. To measure the concentration of the protein, 25 uL of the cell lysate was taken to measure total protein concentration using Pierce™ BCA Protein Assay Kit (Thermo Scientific, 23250).

#### Western Blotting

Protein samples were standardized to 1 ug total protein with DTT (final concentration 0.05M, BIO-RAD, 1610611) and 1X Pierce™ Lane Marker Non-reducing Sample Buffer (Thermofisher 39001). Samples were then boiled at 95°C for 5 minutes and resolved by Mini-PROTEAN® TGX™ Precast 4-15% SDS-PAGE gels. Gels were run for approximately 20 minutes at 120 V. Gels were UV-activated for 1–2 minutes prior to transfer. Proteins were transferred onto methanol-equilibrated PVDF membranes using the BioRad Trans-Blot Turbo transfer system with standard mixed molecular weight mini-gel settings and 1X Trans-Blot Turbo Transfer Buffer. Successful protein transfer was confirmed via Ponceau S staining (Thermo Scientific, A40000278), after which membranes were washed three times with 1X TBST. Anti-FLAG-HRP conjugated (ABCAM ab49763) and anti-RNA polymerase II CTD repeat YSPTSPS (AbCam ab26721) were diluted 1:5000 with BioRad EveryBlot Blocking buffer (12010020). Membranes were incubated overnight at 4°C with primary antibodies. HRP Goat anti-Rabbit IgG (H+L) Secondary antibody (Invitrogen, 31460) was diluted 1:10,000 in BioRad EveryBlot Blocking buffer and incubated for 30 minutes. Membranes were then washed no less than 5 times in 1X TBST. Then each membrane was incubated for exactly one minute with SuperSignal™ West Femto Maximum Sensitivity Substrate (Thermo Scientific, 34095) and imaged with the BioRAd ChemiDoc MP Imaging System (BIO-RAD).

#### Selection of predicted novel RBP IDR activator/nonactivators

For Figure 4D, *metapredict* (refer to **Prediction of intrinsically disordered regions of RBPs**) was first used to predict the IDRs of all human RBPs^2^ (Gerstberger, 2014). The IDRs sequences were then used as input to run NARDINI+ (refer to **Molecular grammar analysis of intrinsically disordered regions**). The scatter plot was then generated using the z score of Frac Aromatic and Frac K+R from the NARDINI+ output. Novel activators were predicted with z score Frac Aromatic > 0 and z score Frac K+R < 0, and novel non-activators were predicted with z score Frac Aromatic < 0 and z score Frac K+R > 0. More filters were applied to the candidates selected for luciferase assay. For novel activator candidates, they all have z score Frac Aromatic > 2 and z score Frac K+R < -1. Minimum length of 30aa, nuclear localization, and indication of their functions in transcription and RNA processing are required for both novel activators and novel non-activators.

### Synthesis and characterization of SNAP-TMP

#### General information

All commercially available compounds were purchased from Chemscene, Ambeed, TCI, Acros organic, Alfa-Aesar, Oakwood Chemical and Thermo Fisher Scientific. All the solvents and all the reagents were directly used from purchased without any further purification.

#### Instrumentation

Thin-layer chromatography (TLC) was performed on Sorbent Technologies silica plates. Automated flash column chromatography was performed using RediSep Rf silica Gel or C18 column on CombiFlash Rf200 system with internal UV detector. The instrument is available from Teledyne Isco, Inc., NE., USA. Proton nuclear magnetic resonance spectroscopy (1H NMR) and Carbon nuclear magnetic resonance spectroscopy (13C NMR) spectra were recorded on a Bruker NEO 600 MHz NMR and processed by MestReNova. Low-resolution mass spectra were obtained using Liquid-Chromatography-Mass-Spectrometry (LCMS) on Waters instrument, electrospray ionization in either positive or negative mode. High-resolution mass spectra (HRMS) were obtained at the University of Pennsylvania’s Mass Spectrometry Service Center on Waters LC-TOF mass spectrometer (model LCT-XE Primer) using electrospray ionization in positive or negative mode, depending on the analytes. HRMS data analysis was performed using the automated Waters software. Except for compound 2, HRMS data for 1 and 4 is collected in a Bruker scimaX instrument with mass accuracy (internal) up to 600 ppb.

#### Synthesis (Figure S7B)

tert-butyl (4-(((2-amino-6-chloropyrimidin-4-yl)oxy)methyl)benzyl)carbamate SNAP-NBoc **1**. 4-(Boc-aminomethyl)benzyl alcohol (96.9 mg, 0.41 mmol) was dissolved in 5 mL 1:1 THF/DMF cooling to 0°C, followed by slow addition of sodium hydride (18.0 mg, 0.45 mmol, 1.1 equiv.). The reaction mixture was stirred for 30 min followed by addition of 2-amino-4,6-dichloropyridine (67.2 mg, 0.41 mmol, 1 equiv.). The reaction was warmed to room temperature and left stirring overnight. The crude reaction was quenched by sat. NH4Cl and diluted with 20 mL water followed by twice ethyl acetate (10 mL) wash. The organic phase was washed with brine and dried with Na2SO4. and then concentrated in vacuo. The crude mixture was subjected to silica chromatography with a gradient of 5-10% methanol in DCM, yielding 106.0 mg of **1** as white solid (71%). ^1^H NMR (600 MHz, CDCl_3_) δ 7.37 – 7.34 (m, 2H), 7.29 (d, *J* = 7.8 Hz, 2H), 6.15 (s, 1H), 5.30 (s, 2H), 5.16 (s, 2H), 4.87 (t, *J* = 6.2 Hz, 1H), 4.32 (d, *J* = 6.0 Hz, 2H), 1.46 (s, 9H) (Figure S8A).

^13^C NMR (151 MHz, CDCl_3_) δ 171.01, 162.31, 161.08, 156.02, 139.28, 135.22, 128.56, 127.81, 97.47, 79.73, 68.15, 44.51, 28.54 (Figure S8B). **HRMS** (ESI, m/z): C17H22ClN4O3 [M+H]+: 365.137495; found: 365.137383.

4-((4-(((2-amino-6-chloropyrimidin-4-yl)oxy)methyl)benzyl)amino)-4-oxobutanoic acid **2**. SNAP-NBoc **1** (0.5076g, 1.4 mmol) and succinic anhydride (0.1412 g, 1.4 mmol, 1 equiv.) was dissolved in 4 mL DMF followed by the addition of DIEA (1.2 mL, 7 mmol, 5 equiv.) and DMAP (17.0 mg, 0.14 mmol, 0.1 equiv.). The reaction mixture was stirred overnight and aq. HCl was added to adjust pH to 2-3. The precipitated solid was filtered out **2** as 0.3273 g white solid (64% yield). ^1^H NMR (600 MHz, DMSO) δ 12.07 (s, 1H), 8.35 (t, *J* = 6.0 Hz, 1H), 7.39 – 7.35 (m, 2H), 7.25 (d, *J* = 8.0 Hz, 2H), 7.10 (s, 2H), 6.13 (s, 1H), 5.29 (s, 2H), 4.25 (d, *J* = 5.9 Hz, 2H), 2.45 (t, *J* = 6.6 Hz, 2H), 2.38 (t, *J* = 6.6 Hz, 2H) (Figure S8C). ^13^C NMR (151 MHz, DMSO) δ 173.86, 170.98, 170.31, 162.78, 160.01, 139.61, 134.60, 128.33, 127.22, 94.41, 67.25, 41.81, 29.96, 29.10 (Figure S8D). **HRMS** (ESI, m/z): C16H17ClN4O4 [M+H]+: 365.1017; found: 365.1036. Compound **3** was prepared according to the literature^129^. N1-(4-(((2-amino-6-chloropyrimidin-4-yl)oxy)methyl)benzyl)-N4-(3-(4-((2,4-diaminopyrimidin-5-yl)methyl)-2,6-dimethoxyphenoxy)propyl)succinimide **4** SNAP-TMP. TMP-C3-NHBoc **3** (60.0 mg, 0.14 mmol) was stirred in 1 mL TFA/DCM 1:1 solution for 30 min and concentrated in vacuo to remove TFA. Then the resultant oil, CP-COOH **2** (50.4 mg, 0.14 mmol), PyBOP (75.5 mg, 0.14 mmol) was dissolved in 2 mL DMF with 240 uL DIEA. The reaction mixture was stirred for 30 min and quenched with 10 mL water followed by twice extraction with ethyl acetate (8 mL). The organic phase was washed with brine, dried with Na2SO4. and then concentrated in vacuo. The crude mixture was subjected to column chromatography with a gradient of 10-20% methanol in DCM, yielding 48.1 mg of **4** as light yellow solid (51% yield). ^1^H NMR (600 MHz, DMSO) δ 8.34 (t, *J* = 6.0 Hz, 1H), 7.79 (t, *J* = 5.6 Hz, 1H), 7.51 (s, 1H), 7.40 – 7.34 (m, 2H), 7.28 – 7.21 (m, 2H), 7.10 (s, 2H), 6.55 (s, 2H), 6.25 (s, 2H), 6.12 (s, 1H), 5.85 (s, 2H), 5.28 (s, 2H), 4.24 (d, *J* = 5.9 Hz, 2H), 3.81 (t, *J* = 6.3 Hz, 2H), 3.71 (s, 6H), 3.53 (s, 2H), 3.19 (q, *J* = 6.6 Hz, 2H), 2.41 – 2.28 (m, 4H), 1.70 (p, *J* = 6.6 Hz, 2H) (Figure S8E). ^13^C NMR (151 MHz, DMSO) δ 171.86, 171.62, 170.78, 163.26, 162.82, 162.02, 160.48, 154.79, 153.30, 140.12, 136.01, 135.17, 135.06, 128.82, 127.68, 106.53, 106.31, 94.89, 70.93, 67.74, 56.32, 49.07, 42.27, 40.53, 40.41, 40.27, 40.13, 39.99, 39.85, 39.71, 39.58, 36.38, 33.39, 31.29, 30.27 (Figure S8F). **HRMS** (ESI, m/z): C32H39ClN9O6 [M+H]+:680.270634; found:680.270386

### Statistical analysis

For Figure 1D, Figure S1C-S1E, Figure 4C and 4E, Figure 5A, and Figure S7A, ordinary one-way ANOVA in Prism was used for statistical analysis. All data were compared to the “Gal4-alone” condition. For Figure S4B, p-value was calculated with Mann-Whitney U test. All statistical information is provided in Table S1.

## Acknowledgements

We thank Dr. Huaiying Zhang, Dr. Biplab KC, and Dr. Joel McManus from Carnegie Mellon University for helpful discussion, reagents, and consultation regarding the synthetic condensate experiments. We would also like to thank the ENCODE consortium for providing ChIP-seq data, specifically from the laboratories of Dr. Xiang-Dong Fu and Dr. Gene Yeo.

## Author contributions

J.E.H. and H.W. conceptualized the project. H.W., B.K.W., G.M., D.X., E.W., D.M.-F., L.P., and C.D. performed the experiments. H.W., G.M., S.S., J.E.H., and A.B. performed data analysis. J.E.H. and H.W. wrote the original draft with the help from all authors. J.E.H and D.M.C supervised and supported the study.

## Materials & Correspondence

Correspondence and material requests should be addressed to.

## Figure Legends

**Figure S1:**
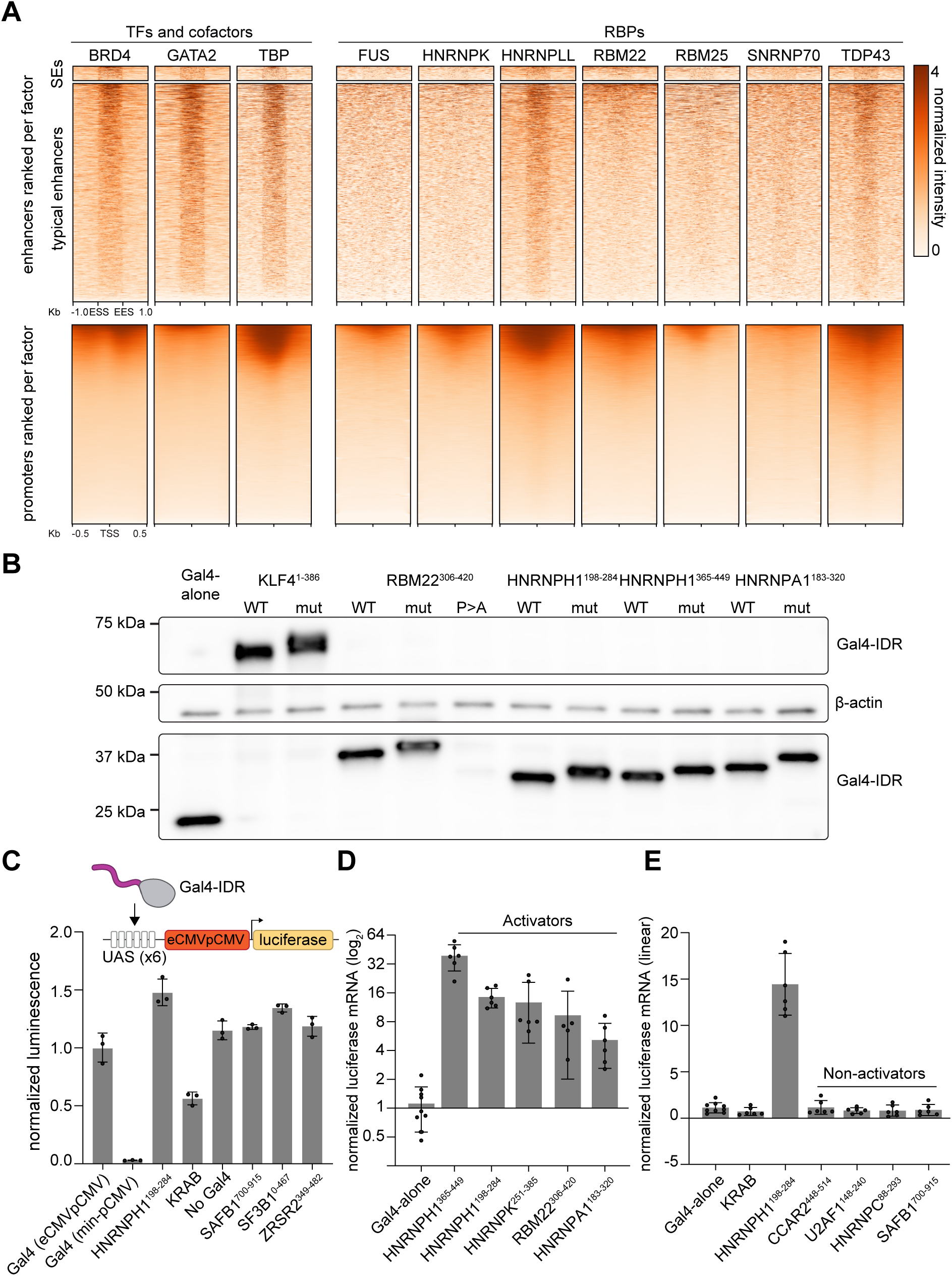
RBP genomic enrichment and reporter assays. A. ChIP–seq heatmap of selected RBPs at super-enhancers, enhancers and promoters. B. Western blot showing the WT and mutant Gal4-IDR protein expression levels C. Bar plots showing the normalized luminescence of Gal4-RBP IDR luciferase assay using a luciferase plasmid with a strong promoter. (statistics in Table S1) D. Bar plot depicting the RT-qPCR validation of select RBP IDR activators in Gal4-RBP IDR luciferase assay (statistics in Table S1). E. Bar plot depicting the RT-qPCR validation of select RBP IDR non-activators in Gal4-RBP IDR luciferase assay (statistics in Table S1).

**Figure S2:**
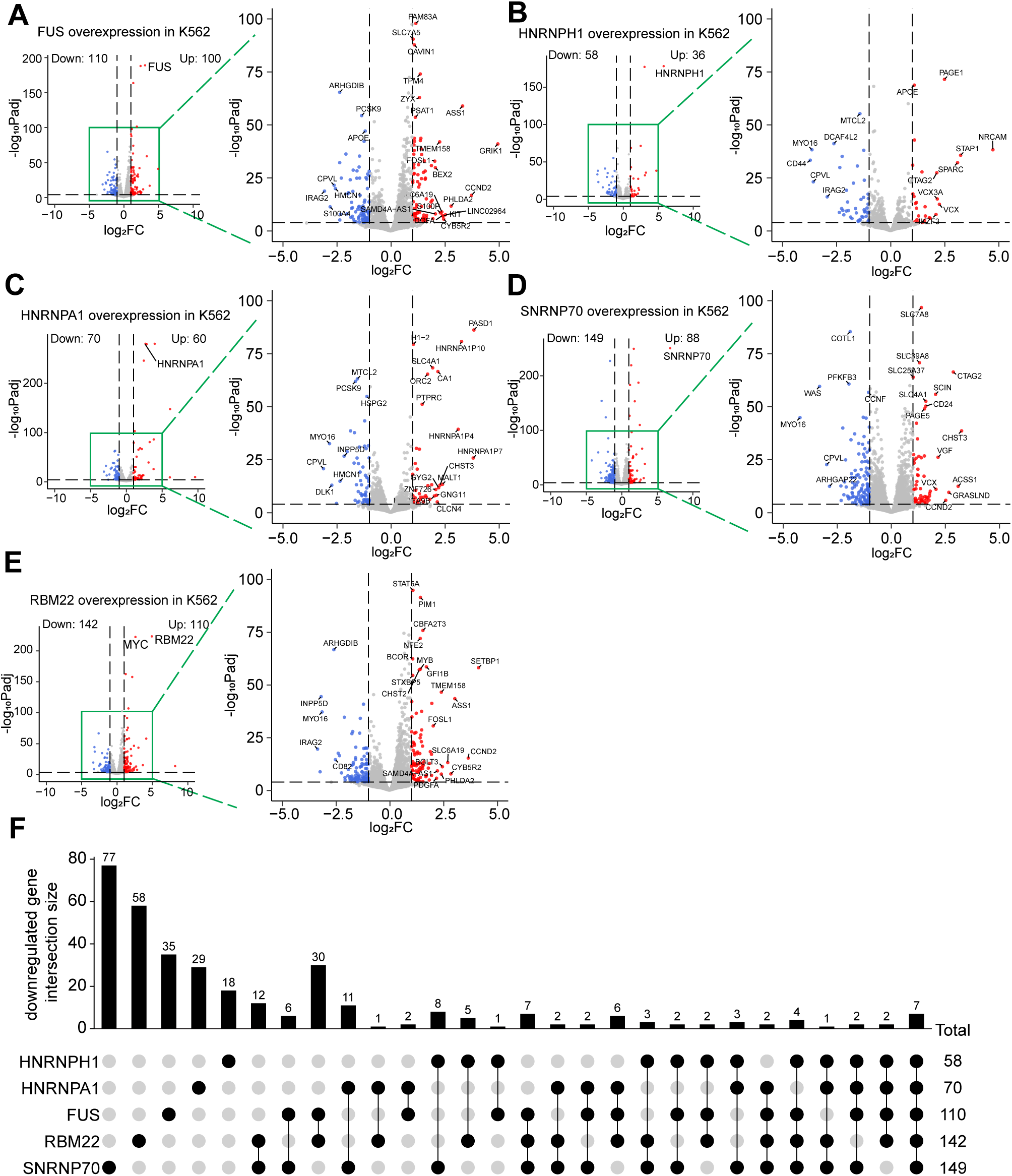
Genes regulated by RBP activators. A-E. Volcano plots of RNA-seq in doxycycline-inducible RBP overexpression K562 cell lines. Significant downregulated genes (padj < 0.05, FC < -1) are highlighted in blue and significant upregulated genes (padj < 0.05, FC > 1) are highlighted in red. F. Upset plot showing the downregulated gene intersection between 5 RBPs.

**Figure S3:**
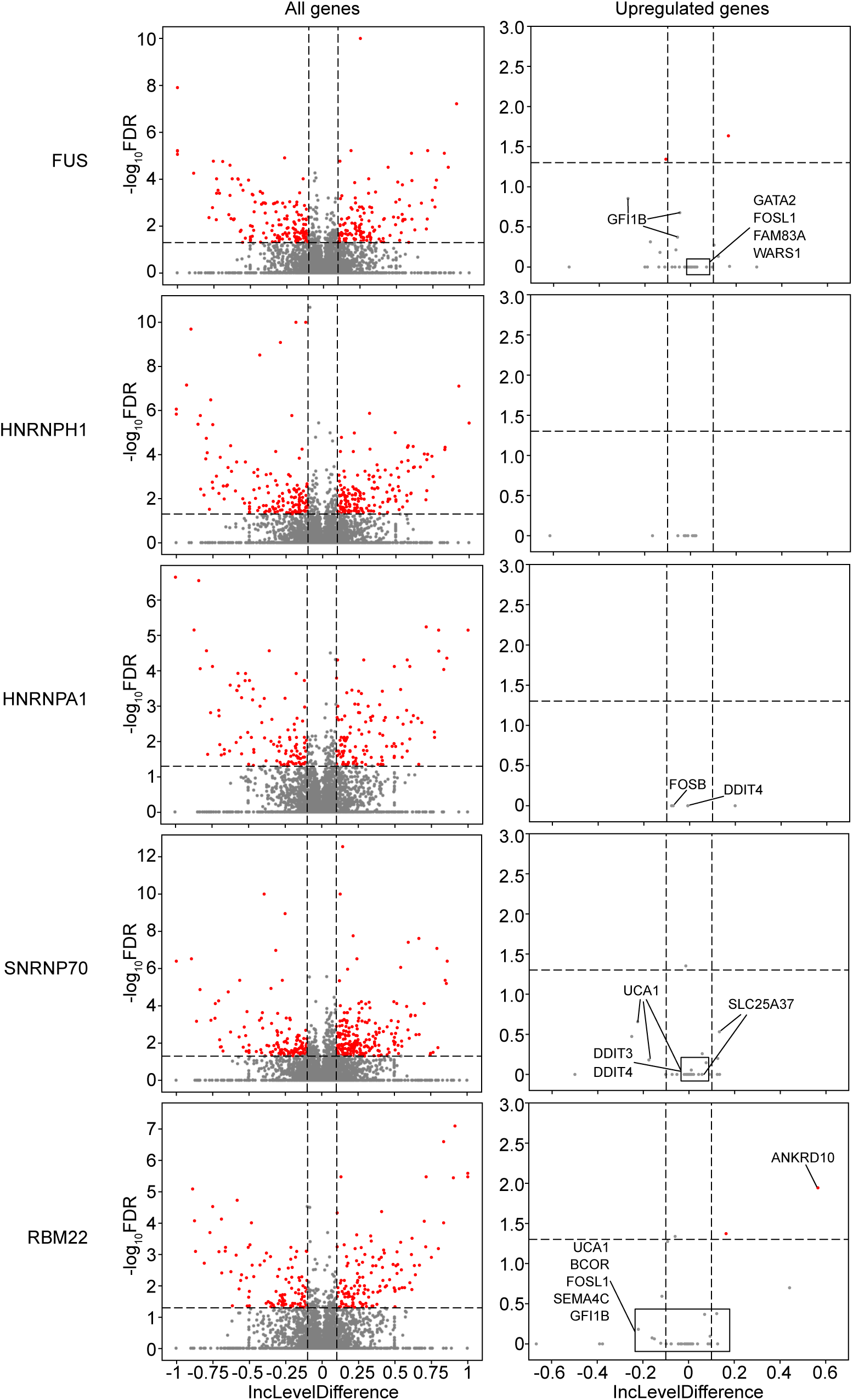
Changes in splicing during acute RBP expression. Left lane: Volcano plots of an example of alternative splicing (A3SS) at all genes upon RBP overexpression. Right lane: Volcano plots of alternative splicing (A3SS) at upregulated super-enhancer-associated genes upon RBP overexpression (significant alternative splicing defined as |IncLevelDifference| > 0.1, FDR < 0.05).

**Figure S4:**
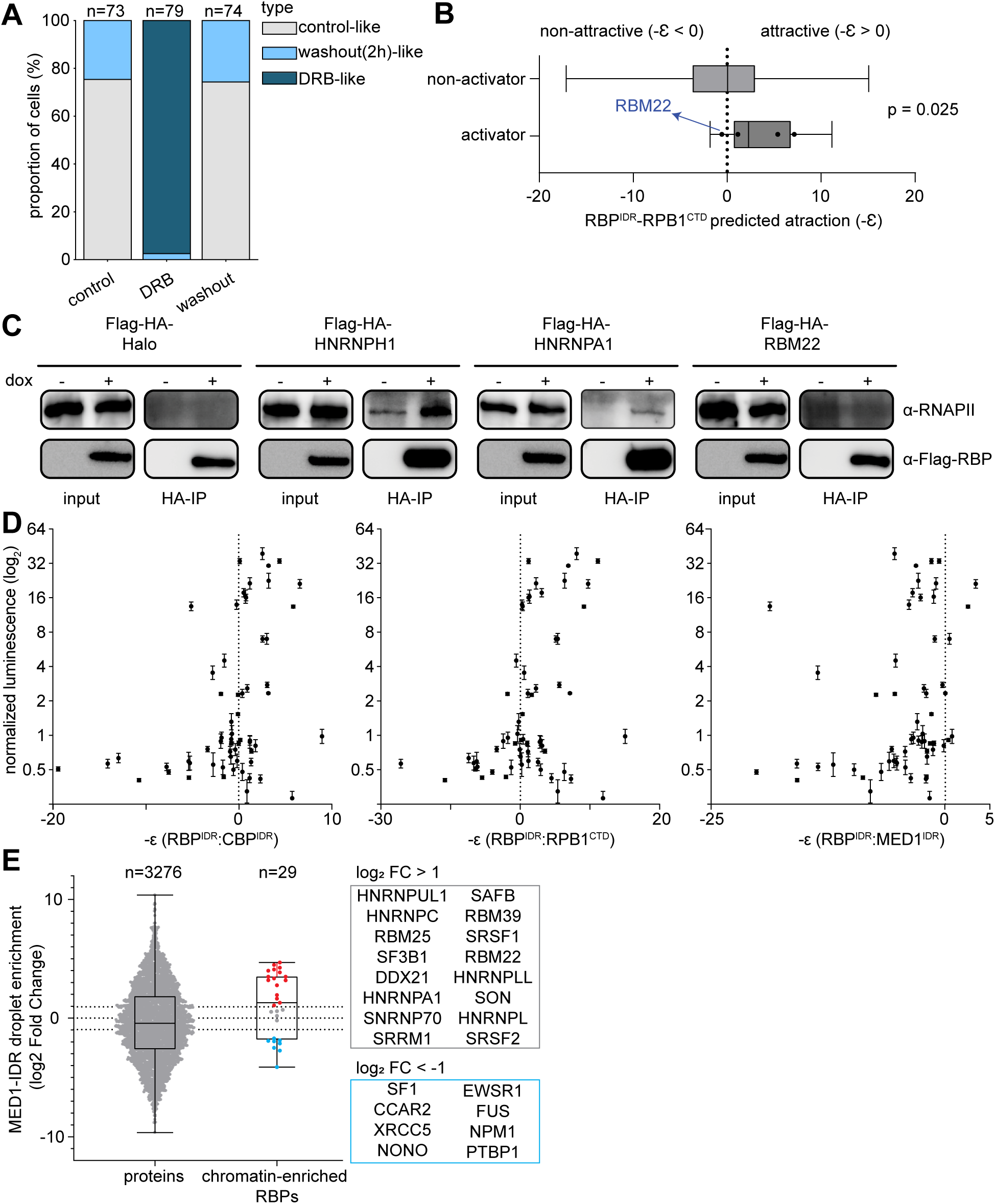
RNA and protein interactions for RBP activators. **A.** Scoring of Hnrnph1 nuclear localization pattern (p value: DRB(100μM) vs. DRB-washout: 6.71e-32; DRB(100μM) vs. clt: 7.25e-32; DRB-washout vs. ctl: 1; pairwise-Chi-square test with Holm correction). B. Box plot showing the prediction interaction between tested RBP IDRs with the C-terminal domain of RPB1 using FINCHES. (p = 0.025; Mann-Whitney U test) C. Co-IP of RBP with RNAPII in doxycycline-inducible RBP overexpression K562 cell lines. D. Scatter plot showing the relationship between normalized luminescence from the Gal4–luciferase assay and FINCHES-predicted interactions of RBP IDRs with MED1 IDR, BRD4 IDR, and the RPB1 CTD. E. Box plot showing the enrichment of select RBPs in MED1-IDR condensate. Data from Lyons et al., 2023.

**Figure S5:**
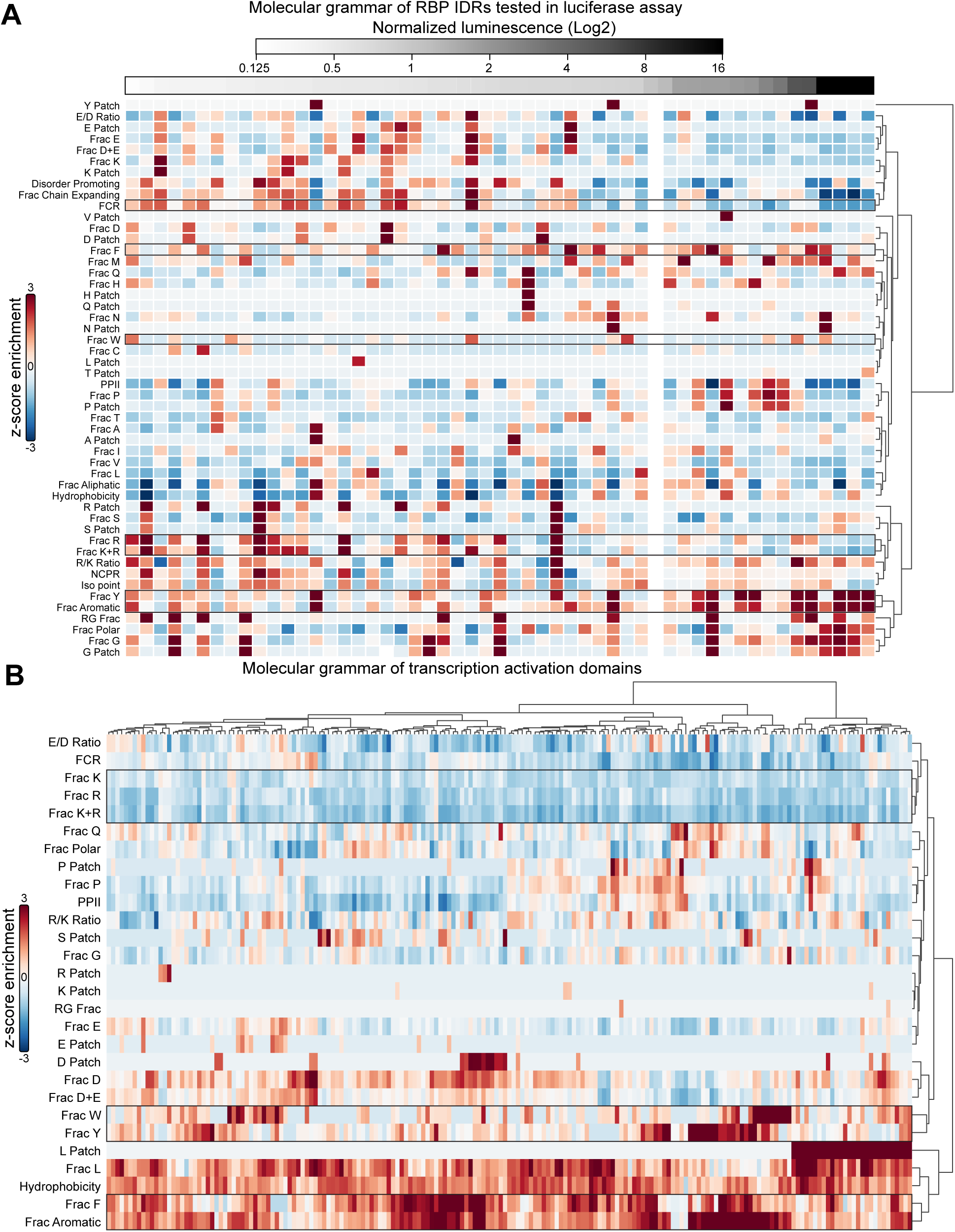
Feature enrichments for RBPs and TFs. **A.** NARDINI+ analysis heatmap of amino acid composition of IDR tested in the Gal4 luciferase assay (ranked by their normalized luminescence) B. NARDINI+ analysis heatmap of amino acid composition of transcription activation domains from DelRosso et al., 2023.

**Figure S6:**
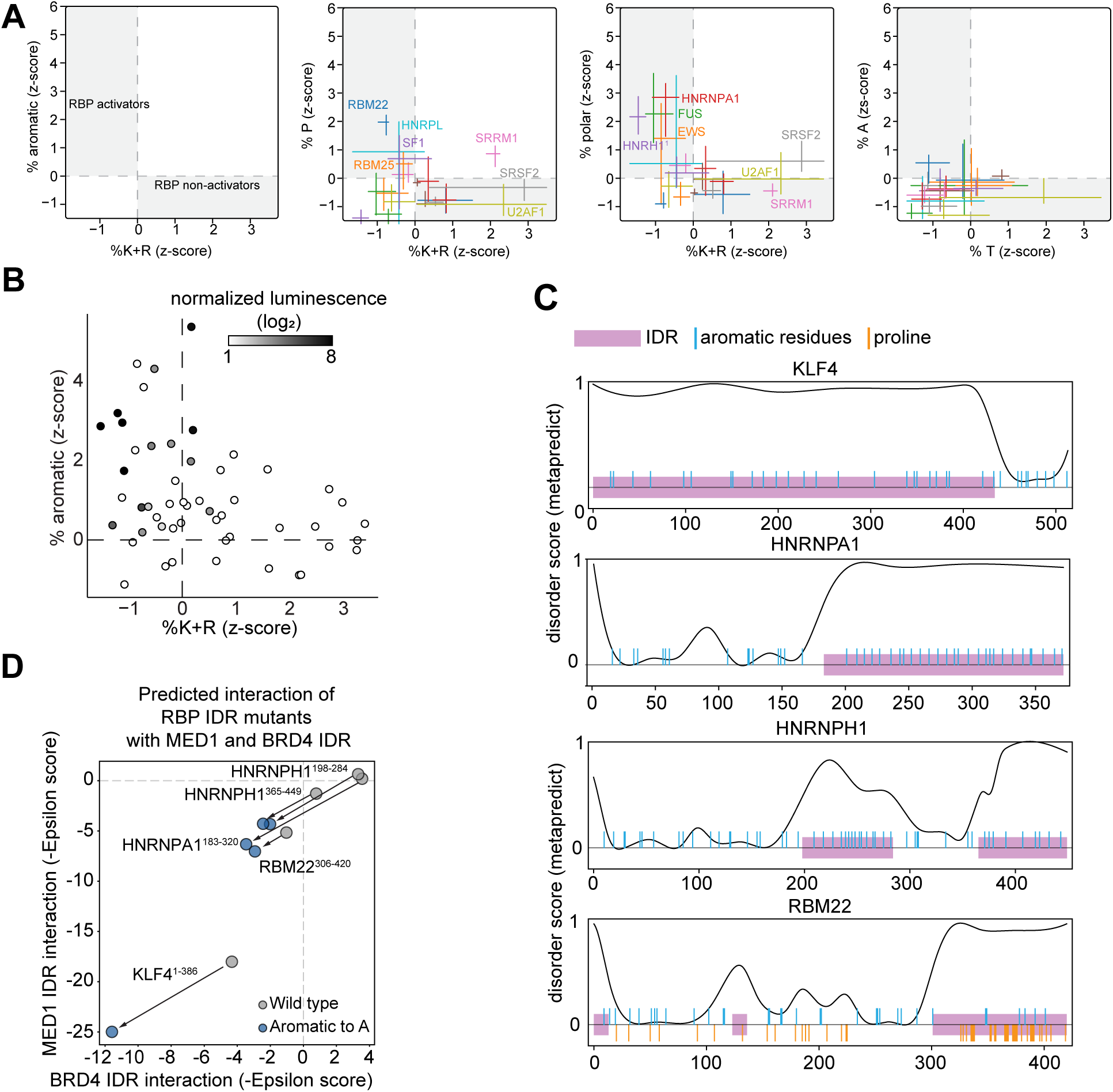
Conservation and perturbation of RBP molecular grammar. **A.** NARDINI+ analysis of the enrichment of other amino acid composition features of RBP orthologues across vertebrates. B. Scatter plot showing the enrichment of aromatic and basic residues of RBP IDRs tested in Gal4 luciferase assay with NARDINI+ analysis. Each IDR is colored based on its normalized luminescence values in the assay. C. Depiction of the IDR regions of select RBPs with positions of aromatic and proline residues labeled. D. Scatter plot of predicted interaction between RBP IDR mutants with MED1 and BRD4 by FINCHES analysis

**Figure S7:**
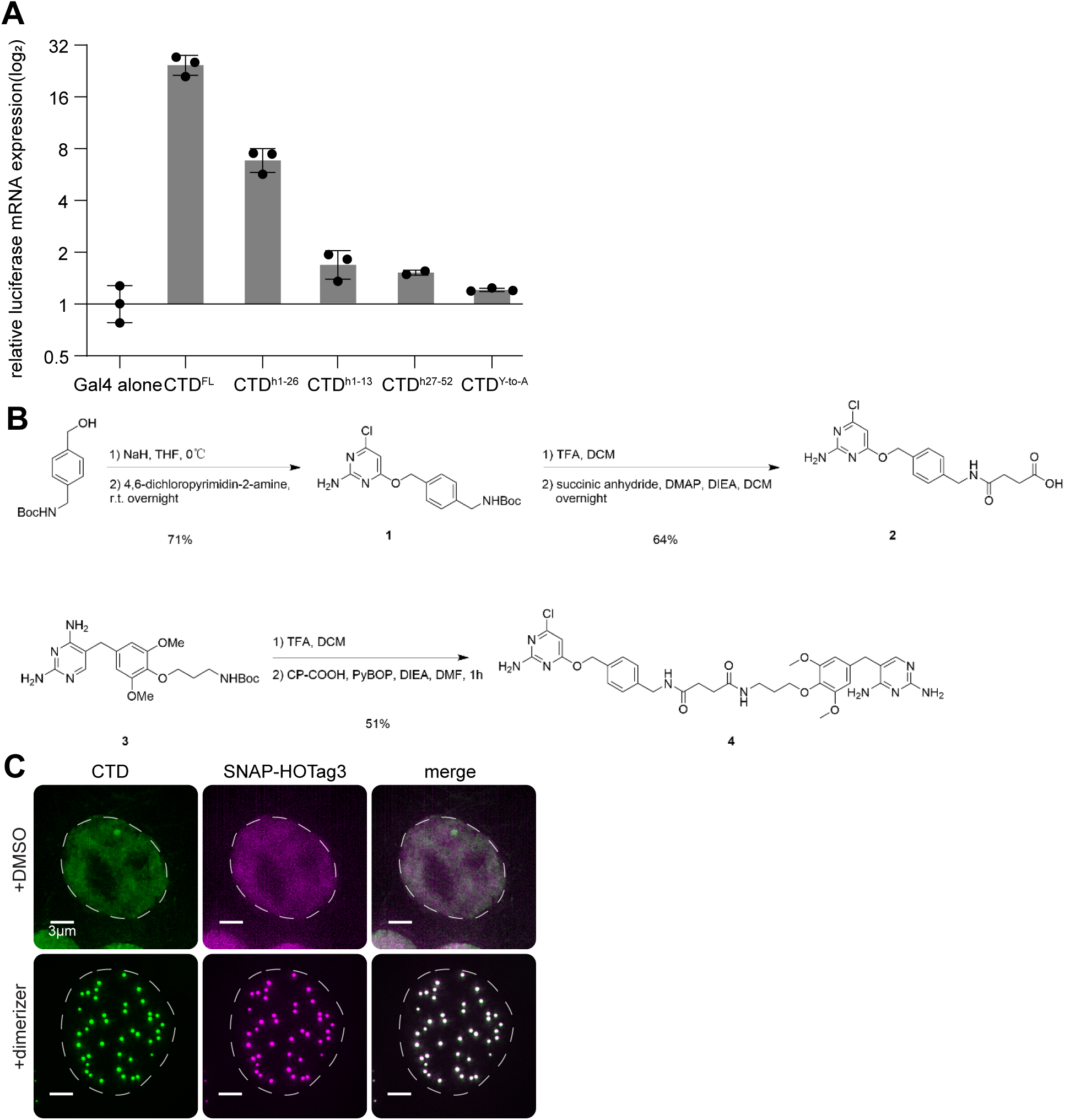
CTD reporter assays and scaffold-based condensate formation. **A.** Bar plot depicting the RT-qPCR validation of RPB1 CTD variants in Gal4-RBP IDR luciferase assay (statistics in Table S1). B. Synthetic scheme of SNAP-TMP (ST) C. Max projection of selected z slices images of single nuclei expressing CTD-mStayGold-eDHFR and SNAP-HOTag3 in the presence and absence of dimerizer.

**Figure S8:**
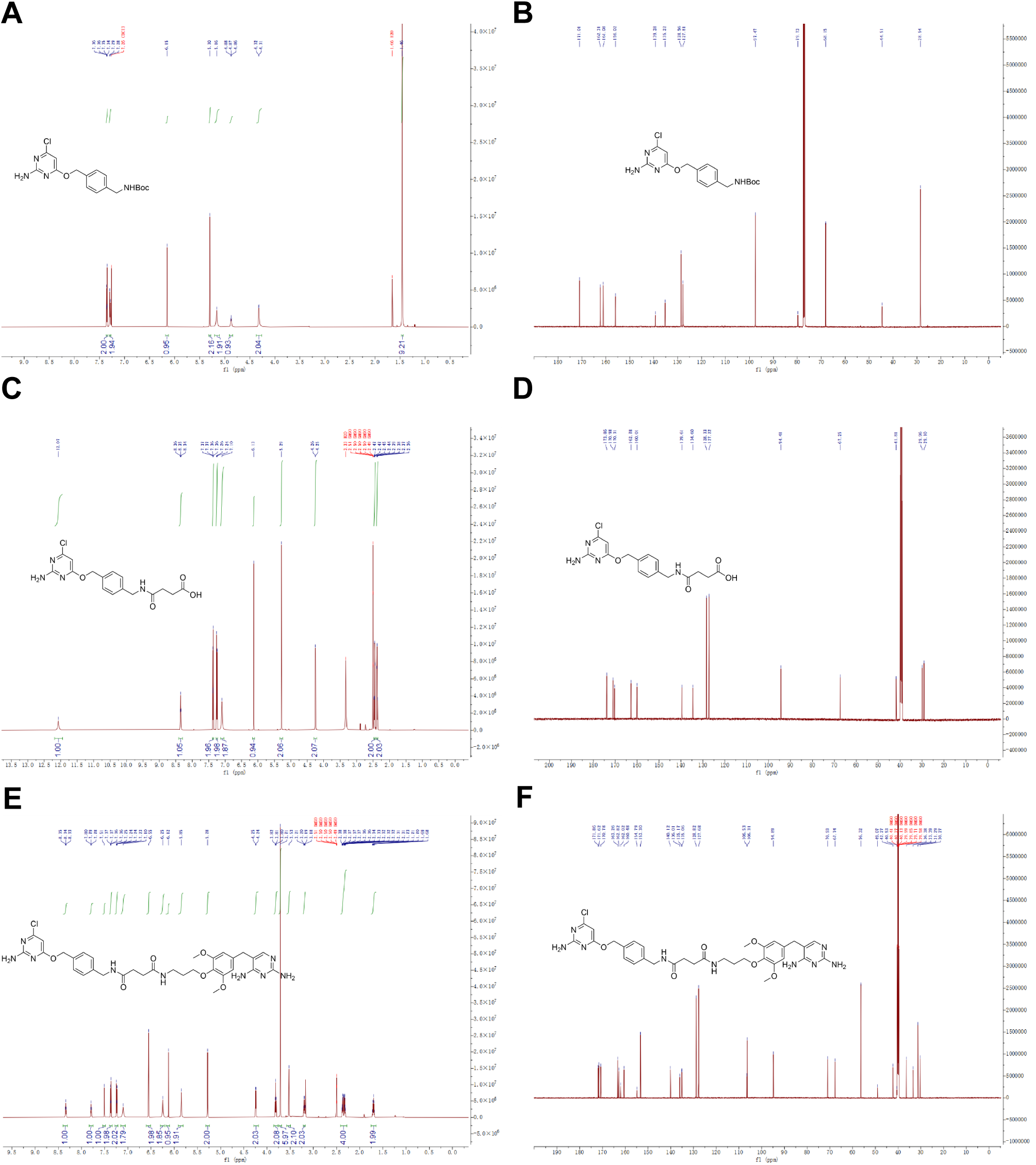
NMR spectra of each product during SNAP-TMP synthesis. **A.** ¹H NMR spectrum of **1** in CDCl3 (600 MHz) B. 13C NMR spectrum of **1** in CDCl3(151 MHz) C. ¹H NMR spectrum of **2** in DMSO-d6 (600 MHz) D. ¹³C NMR spectrum of **2** in DMSO-d6 (151 MHz) E. ¹H NMR spectrum of **4** in DMSO-d6 (600 MHz) F. ¹³C NMR spectrum of **4** in DMSO-d6 (151 MHz)

Table S1 Statistical analysis of luciferase assays and RT-qPCR

Table S2 DESeq2 output of K562 RNA-seq data in **Figure 2**

Table S3 The amino acid composition feature z-scores of all RBP IDRs using NARDINI+

Table S4 Plasmids and oligos used in this study

Movie S1 -

Time-lapse imaging of a HA-mScarlet3-Med1 HA-mStayGold-Rbm22 mouse embryonic stem cell. Left: merged videos (Med1: green; Rbm22: magenta); right: Heatmaps showing the colocalization dynamics between Rbm22 and Med1.

Movie S2 -

Time-lapse imaging of a HA-mScarlet3-Med1 HA-mStayGold-Hnrnph1 mouse embryonic stem cell. Left: merged videos (Med1: green; Hnrnph1: magenta); right: Heatmaps showing the colocalization dynamics between Hnrnph1 and Med1.

